# Schizophrenia-associated 22q11.2 deletion elevates striatal acetylcholine and disrupts thalamostriatal projections to produce amotivation in mice

**DOI:** 10.1101/2025.06.20.660782

**Authors:** Mary H. Patton, Brett J.W. Teubner, Kristen T. Thomas, Sharon L. Freshour, Alexandra J. Trevisan, Andrew B. Schild, Cody A. Ramirez, Jay B. Bikoff, Stanislav S. Zakharenko

## Abstract

Schizophrenia is a complex neurodevelopmental disorder characterized by cognitive dysfunction, hallucinations, and negative symptoms such as amotivation. Negative symptoms are largely resistant to current antipsychotic treatments, and the neural circuits underlying amotivational states remain poorly defined. Here, using a mouse model of schizophrenia-associated 22q11.2 deletion syndrome (22q11DS), we report amotivation and weakened glutamatergic synaptic transmission between the thalamic parafascicular nucleus (Pf) and the dorsomedial striatum (DMS). Thalamostriatal disruption is attributed to hyperactivity of striatal cholinergic interneurons (CHIs), which is associated with enhanced *Trpc3* and *Pex51* (*Trip8b*) gene expression. Elevated acetylcholine levels in the DMS act on presynaptic M2 muscarinic receptors to weaken Pf-DMS glutamatergic transmission. Importantly, disruption of Pf-DMS synaptic transmission or hyperactivation of CHIs are each sufficient to cause amotivation in wild-type mice. These results identify a striatal hypercholinergic state and subsequent thalamostriatal disruption as core pathogenic events causing amotivation in 22q11DS, providing potential therapeutic targets.

## Introduction

Schizophrenia (SCZ) is a neurodevelopmental disorder characterized by a collection of positive, negative, and cognitive symptoms. Unlike positive symptoms, negative symptoms are particularly resistant to current antipsychotic treatment options, representing an unresolved therapeutic target in SCZ.^1–7^ One negative symptom is a decreased motivational drive, or amotivation,^8–15^ which is associated with poor functional outcomes in patients.^16–20^ The development of effective treatments for amotivation remains limited due to a lack of mechanistic knowledge of the neural circuits underlying native motivation.

Appetitive drive in reward-seeking arises from dopaminergic signaling within the nucleus accumbens.^21–26^ However, nucleus accumbens signaling is typically involved in drug-selective motivational behaviors rather than the natural reward-seeking behaviors that are disrupted in SCZ,^27–37^ with exceptions.^38^ Therefore, disruptions in distinct neural circuitry likely underlie amotivation. The dorsomedial striatum (DMS) governs motivated, goal-directed behavior by integrating sensorimotor and cognitive information.^39–44^ Abnormalities in dorsal striatal functioning are linked to negative SCZ symptoms such as amotivation.^45–48^

Given the high degree of inhibition within the DMS, the excitatory drive of principal medium spiny GABAergic projection neurons (MSNs) must be substantial for these neurons to fire and encode a behavioral response. Information arising from thalamic and cortical inputs is integrated in the DMS at the cellular level by convergence onto MSNs.^49–57^ MSN output is also modulated through intrastriatal dopaminergic and cholinergic signaling. Dopamine release in the DMS is inducible through the activation of the parafascicular nucleus of the thalamus (Pf) and is regulated through acetylcholine (ACh) signaling from local cholinergic interneurons (CHIs).^58–62^ Despite a strong link between disrupted ACh signaling and SCZ,^63,64^ the exact involvement of the Pf-MSN-CHI trisynaptic striatal circuitry in motivated behavior remains unknown.

Here, we investigated the neural circuits underlying amotivation in a mouse model of 22q11 deletion syndrome (22q11DS), one of the strongest genetic predictors of SCZ.^65–72^ We show that amotivation in 22q11DS mice is caused by weakened glutamatergic synaptic transmission from thalamic Pf input to MSNs of the DMS. This thalamostriatal disruption is consistent with previous findings implicating the thalamus in SCZ.^73^ We show that CHI hyperactivity mediates thalamostriatal disruption in 22q11DS mice by producing a hypercholinergic state. We also present single-nucleus RNA-sequencing (snRNA-seq) data that identify possible mechanisms of increased CHI activity.

## Results

### 22q11DS mice demonstrate amotivation

To determine whether 22q11DS mice have motivation deficits, we used the *Df(16)1/+* mouse model, which has been shown to phenocopy many of the behavioral aspects of 22q11DS (**Figure 1A**).^74–77^ We tested adult female and male *Df(16)1/+* and wild-type (WT) littermates on the progressive ratio (PR) task, an operant reward-seeking task that is highly sensitive to differences in motivation levels^19,78,79^ (**Figure 1B**). *Df(16)1/+* mice lever pressed fewer times and earned fewer rewards in both the PR+1 and PR+4 tasks than WT littermates (**Figure 1C, D, Figure S1A**). This difference in lever pressing was specific to the PR task, as performance on the random ratio (RR) task was similar between genotypes (**Figure S1B**). To determine whether *Df(16)1/+* mice show a bias toward low-effort food reward when compared to WT littermates, we next tested the mice on a low-effort choice task in which they chose between a fixed ratio 1 (FR1) schedule to obtain sweetened condensed milk (SCM) and freely available food chow (FR1 choice task). There was no difference between genotypes in lever presses (**Figure 1E**) or the amount of chow consumed (**Figure S1C**) on this task.

**Figure 1.**
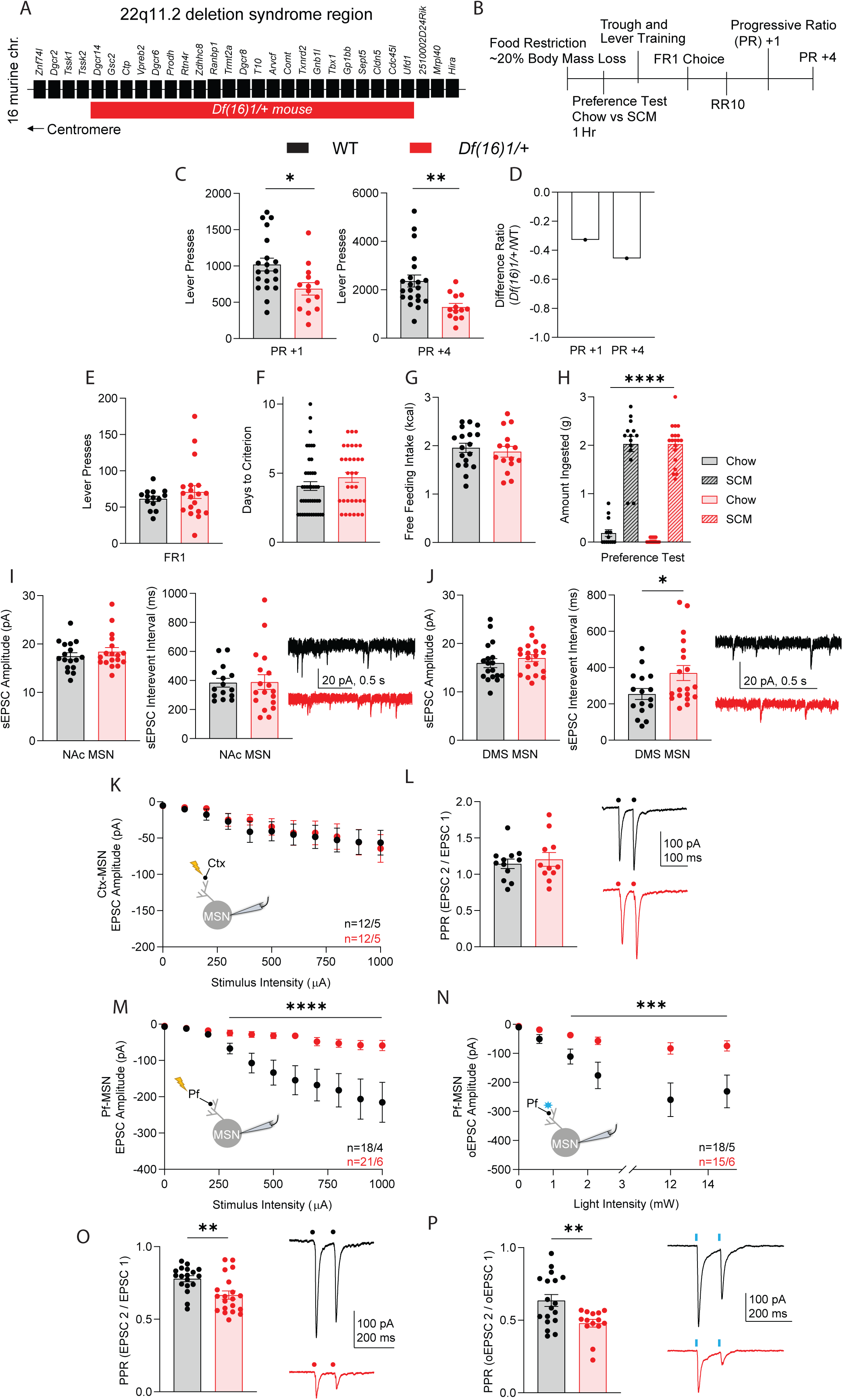
22q11DS mice show amotivation and weakened synaptic transmission at the parafascicular nucleus (Pf) to medium spiny neuron (MSN) synapse. (A) Diagram depicting the genes in the 22q11.2 syntenic region of mouse chromosome 16. Individual genes are represented by rectangles. Horizontal red bar represents the region of the hemizygous deletion in the *Df(16)1/+* mouse model of 22q11 deletion syndrome (22q11DS). (B) Timeline of training and reinforcement schedules for behavioral studies. (C) Left: In the progressive ratio (PR) +1 task, *Df(16)1/+* mice (red) lever pressed fewer times than WT littermates (black; two-way ANOVA, \**p* = 0.02). Right: In the PR+4 task, *Df(16)1/+* mice lever pressed fewer times than WT littermates (two-way ANOVA, \*\**p* = 0.004). (D) The average ratio of the difference in performance between *Df(16)1/+* mice and WT littermates was –0.33 for PR+1 and –0.46 for PR+4. (E) There was no difference in lever presses on the fixed ratio 1 (FR1) task between genotypes (two-way ANOVA, *p* = 0.37). (F) The average number of training days needed to reach criterion to progress to the operant tasks was similar between genotypes (two-way ANOVA, *p* = 0.2). (G) Free feeding intake of sweetened condensed milk (SCM) was similar between genotypes (two-way ANOVA, *p* = 0.62). (H) Both WT and *Df(16)1/+* mice preferred SCM over standard rodent chow when both were freely available (mixed-design ANOVA, \*\*\*\**p* < 0.0001). There was no difference in the amount of SCM consumed between genotypes (Tukey’s multiple comparisons, *p* > 0.99). The animal’s sex significantly affected food preference (mixed-design ANOVA, \**p* = 0.048). (I) There was no difference in the amplitude (left; unpaired *t* test, *p* = 0.41) or interevent interval (center; unpaired *t* test, *p* = 0.95) of spontaneously occurring excitatory postsynaptic currents (sEPSCs) in nucleus accumbens (NAc) medium spiny neurons (MSNs) between genotypes. Right: Example traces of sEPSCs from NAc MSNs from WT and *Df(16)1/+* mice. (J) There was no difference in the amplitude of sEPSCs in dorsomedial striatal (DMS) MSNs between genotypes (left; unpaired *t* test, *p* = 0.39). The interevent interval was significantly higher between sEPSC events in MSNs from *Df(16)1/+* mice (center; unpaired *t* test, \**p* = 0.04). Right: Example traces of sEPSCs from DMS MSNs from WT and *Df(16)1/+* mice. (K) Electrically evoked EPSC amplitudes at the cortical (Ctx)-MSN synapse in response to increasing stimulus intensity were similar between genotypes (mixed-design ANOVA, *p* = 0.98). Inset shows the recording configuration. (L) Left: The average paired pulse ratio (PPR) of evoked EPSCs at the Ctx-MSN synapse was similar between genotypes (unpaired *t* test, *p* = 0.6). Right: Example traces of evoked Ctx-MSN EPSCs from WT and *Df(16)1/+* mice. (M) There was a significant decrease in the amplitude of electrically evoked EPSCs at the Pf-MSN synapse in *Df(16)1/+* mice compared to WT littermates (mixed-design ANOVA, \*\*\*\**p* < 0.001). Inset shows the recording configuration. (N) There was a significant decrease in the amplitude of optogenetically evoked EPSCs (oEPSC) at the Pf-MSN synapse in *Df(16)1/+* mice (mixed-design ANOVA, \*\*\**p* < 0.0003). Inset shows the recording configuration. (O) Left: There was a significant difference in the PPR of electrically evoked EPSCs at the Pf-MSN synapse between genotypes (unpaired *t* test, \*\**p* = 0.003). Right: Example traces of electrically evoked Pf-MSN EPSCs from both genotypes. (P) Left: There was a significant difference in the PPR of optogenetically evoked EPSCs at the Pf-MSN synapse between genotypes (unpaired *t* test, \*\**p* = 0.005). Right: Example traces of optogenetically evoked Pf-MSN EPSCs from both genotypes. All data shown are mean ± SEM with individual data points overlaid in (C–J), (L), (O), (P). Unless noted, there were no sex differences. *n* = the number of cells/number of mice. Lightning bolts (K, M) represent electrical stimulation. Circles (L, O) represent stimulus artifacts. Blue star (N) represents optical stimulation. Blue lines (P) represent the optical stimulus. See **Figure S1** for additional behavioral controls and for additional electrophysiology measures.

*Df(16)1/+* and WT littermates also showed a similar ability to learn the operant task (**Figure 1F**). *Df(16)1/+* and WT mice consumed similar amounts of SCM when given free access to it (**Figure 1G**), and both groups significantly preferred SCM over chow to a similar degree in a food preference test (**Figure 1H**). Moreover, we observed no gross locomotor difference between genotypes or differences in the amount of time the mice spent in the center of an open field (**Figure S1D, E**). These findings show that *Df(16)1/+* mice can learn complex operant tasks, do not show anhedonia towards SCM, and do not have gross locomotor disruptions. Taken together, these data demonstrate a specific deficit in motivation in the *Df(16)1/+* mouse model of 22q11DS during a high-demand motivation task.

### 22q11DS mice have weakened glutamatergic synaptic transmission at the Pf-MSN synapse in the DMS

To explore the synaptic mechanisms underlying this amotivation state, we first considered whether *Df(16)1/+* mice exhibit altered synaptic transmission within the nucleus accumbens, a structure well-established to influence motivation.^80^ We used whole-cell patch-clamp electrophysiology in acute mouse brain slices containing the shell of the nucleus accumbens from WT and *Df(16)1/+* mice to measure spontaneously occurring excitatory postsynaptic currents (sEPSCs) onto MSNs. We observed no differences in the frequency or amplitude of sEPSCs between genotypes (**Figure 1I**). Moreover, we found no differences between genotypes in the excitatory drive of nucleus accumbens MSNs (**Figure S1F**). Therefore, we reasoned that the site of aberrant cellular activity underlying amotivation in *Df(16)1/+* mice is located elsewhere.

We next investigated the firing and synaptic properties of MSNs in the nearby DMS, as the DMS is necessary for motivated, goal-directed actions.^39,40^ We observed similar firing and membrane properties of DMS MSNs between genotypes (**Figure S1G–P**). However, measuring the amplitude and frequency of sEPSC events onto DMS MSNs revealed a lower excitatory drive in *Df(16)1/+* MSNs (**Figure 1J**). This unbiased approach identified a greater interevent interval between sEPSC events in *Df(16)1/+* mice than in WT mice and no difference in the amplitude of these events between genotypes. These findings suggest a decrease in the presynaptic release of glutamate onto MSNs. DMS MSNs receive excitatory inputs from two major sources: primary sensorimotor and prefrontal cortices, and the Pf thalamic nucleus.^53,81,82^ We used the input-output relationships between stimulation intensity of each of these regions to investigate the source of the excitatory deficit onto MSNs in oblique horizontal brain slices.^55^ The electrically evoked synaptic transmission and the resulting paired pulse ratio (PPR) from cortical inputs onto MSNs were similar between *Df(16)1/+* mice and WT littermates (**Figure 1K, L**). However, electrically evoking Pf release onto MSNs resulted in weaker synaptic transmission in *Df(16)1/+* mice compared to WT (**Figure 1M**). We also detected this Pf-MSN synaptic transmission deficit by optically activating Pf neurons with the light-sensitive opsin channelrhodopsin (ChR2) (**Figure 1N**). Both electric and optogenetic approaches revealed a difference in PPR at the Pf-MSN synapse between genotypes (**Figure 1O, P**), further implicating the presynaptic locus in the thalamostriatal disruption in 22q11DS mice.

### Dampening Pf-DMS synaptic transmission causes amotivation

To investigate the behavioral relevance of these synaptic deficits, we used a chemogenetic approach (designer receptors exclusively activated by designer drugs [DREADDs]^83,84^) to mimic Pf transmission deficits in WT mice. To specifically target the Pf-DMS pathway, we bilaterally injected *rAAV2-retro*, an engineered AAV2 variant that enables retrograde access to projection neurons,^85^ that contains Cre-recombinase into the DMS. We then injected a Cre-dependent virus encoding the inhibitory DREADD receptor hM4Di bilaterally into the Pf into the animal (**Figure 2A, B**). We tested these mice on operant tasks in the presence and absence of the DREADD agonist compound 21 (C21). The disruption of Pf-DMS synaptic transmission after the injection of C21 produced amotivation: C21-injected mice performed fewer lever presses and earned fewer rewards than vehicle-injected mice in both the PR+1 and the PR+4 tasks (**Figure 2C–E**). The reduction in lever pressing on the PR+4 task by experimentally decreasing Pf-DMS synaptic transmission was smaller (one-sample *t* test with µ = –0.455, *p* = 0.002) than that between *Df(16)1/+* and WTs (**Figure 2D** and **Figure 1D**).

**Figure 2.**
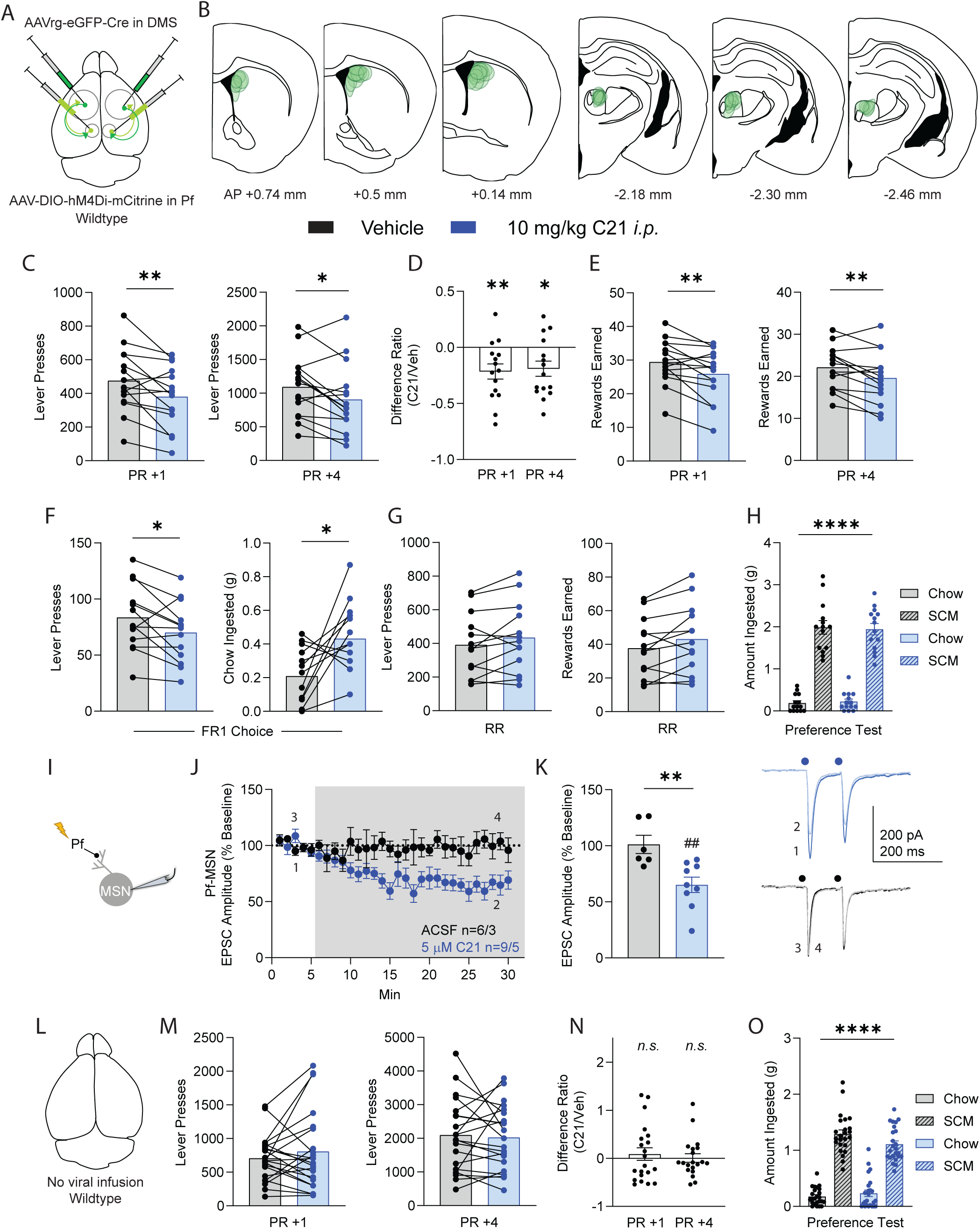
Dampening Pf-DMS synaptic transmission causes amotivation. (A) Schematic of the injection paradigm. (B) Histologic verification of injection sites in the DMS (AP +0.74 to +0.14 mm from bregma) and Pf (AP –2.18 to –2.46). Circles are virus expression from individual mice. (C) Mice injected with the DREADD agonist Compound 21 (C21; blue) lever pressed less than vehicle-injected mice (black) in the PR+1 task (left; two-way repeated measure (RM) ANOVA, \*\**p* = 0.009) and in the PR+4 task (right; two-way RM ANOVA, \**p* = 0.03). (D) The average difference ratio between vehicle and C21 injection was –0.22 ± 0.07 in the PR+1 task performance and –0.19 ± 0.07 in the PR+4 performance. These ratios were significantly different from a theoretical mean of 0 (one-sample *t* test for PR+1, \*\**p* = 0.007; for PR+4, \**p* = 0.01). (E) The number of rewards earned on the PR+1 (left) and PR+4 (right) tasks was lower with C21 injection compared to vehicle (two-way RM ANOVA for PR+1, \*\**p* = 0.006; for PR+4, \*\**p* = 0.009). (F) Left: There were significantly fewer lever presses in the FR1 choice task after C21 injection compared to vehicle (two-way RM ANOVA, \**p* = 0.02). Right: There was significantly higher ingestion of freely available standard rodent chow in the FR1 choice task after C21 injection compared to vehicle (two-way RM ANOVA, \**p* = 0.02). (G) There was no difference in the number of lever presses (left) or the rewards earned (right) on the RR task between C21 and vehicle injection (two-way RM ANOVA for lever presses, *p* = 0.16; for rewards earned, *p* = 0.07). (H) Mice preferred SCM over standard rodent chow when both were freely available regardless of injection type (three-way ANOVA, \*\*\*\**p* < 0.0001). There was no difference in the amount of SCM consumed based on injection type (Tukey’s multiple comparisons, *p* = 0.97). (I) Schematic of the recording configuration. (J) In mice expressing hM4Di in the Pf-DMS pathway, bath application of C21 decreased the evoked amplitude of Pf-MSN EPSCs (blue), whereas bath application of artificial cerebrospinal fluid (ACSF) did not (black). Shaded area depicts the time of drug or ACSF wash. (K) Left: The average Pf-MSN EPSCs in response to ACSF and C21 wash were significantly different (unpaired *t* test, \*\**p* = 0.005). The data analyzed are from the final 5 min of the experiment. The response to ACSF wash was not different from baseline (one-sample *t* test, µ = 100, *p* = 0.89). The response to C21 wash was different from baseline (one-sample *t* test, µ = 100, ##*p* = 0.003). Right, top: Representative traces showing the decrease in amplitude after C21 application (2) compared to baseline (1). Right, bottom: Representative traces showing no difference in EPSC amplitude after ACSF application and at baseline (4 and 3, respectively). The numbers 1–4 correspond to the timepoints labeled in (J). (L) Schematic of the experimental design depicting a WT mouse with no intracranial viral injections. (M) In the absence of DREADDs, injection of C21 did not alter performance on PR+1 (left; two-way RM ANOVA, *p* = 0.33) or PR+4 (right; two-way RM ANOVA, *p* = 0.74). (N) The average difference ratio in performance between vehicle and C21 injection was 0.09 ± 0.13 for PR+1 and 0.006 ± 0.09 for PR+4. These ratios were not different from a theoretical mean of 0 (one-sample *t* test for PR+1, *p* = 0.51; for PR+4, *p* = 0.94). (O) Mice injected with vehicle or C21 preferred SCM over food chow when both were readily available (three-way ANOVA, \*\*\*\**p* < 0.0001). There was no difference in the amount of SCM consumed based on injection type (Tukey’s multiple comparisons, *p* = 0.11). The animal’s sex affected food preference (three-way ANOVA, \**p* = 0.02). All data shown are mean ± SEM with individual data points overlaid in (C–H), (K), (M–O). Unless noted, there were no sex differences. *n* = the number of cells/number of mice. Lightning bolt (I) represents electrical stimulation. Circles (K, right) represent stimulus artifacts. *n.s.*: not significant. See **Figure S2** for additional electrophysiology measures of Pf neurons.

Consistent with the PR data, C21 injection induced a low-effort bias in the FR1 choice task (**Figure 2F**). C21-injected mice lever pressed for SCM less and consumed more freely available chow than did vehicle-injected mice. Interestingly, this low-effort bias was not observed in *Df(16)1/+* mice (**Figure 1E and S1C**). However, consistent with behavior in *Df(16)1/+* mice, lower Pf-DMS synaptic transmission did not alter performance or rewards earned on the RR task (**Figure 2G**), and C21 did not affect the preference for SCM over rodent chow (**Figure 2H**). These findings suggest that amotivation occurs at least partly through deficits in the synaptic transmission at the Pf-DMS circuit.

To verify the function of the hM4Di DREADD used in behavioral studies, we made oblique horizontal slices containing the Pf and DMS of mice injected with Cre-dependent hM4Di and retrogradely-transported Cre-recombinase. The electrically evoked EPSCs at Pf-MSN synapses (**Figure 2I**) were reduced after bath application of 5 µM C21, a concentration comparable to the dose delivered *in vivo* (**Figure 2J, K**). This decrease in synaptic transmission did not occur after bath application of artificial cerebrospinal fluid (ACSF) (**Figure 2J, K**). Moreover, the intraperitoneal injection of C21 to WT mice not expressing DREADD receptors (**Figure 2L**) did not alter performance on the PR+1 or PR+4 task (**Figure 2M, N**). Similarly, C21 injection alone did not alter SCM preference (**Figure 2O**).

### 22q11DS mice have enhanced DMS cholinergic activity

Based on these findings and previous studies showing altered synaptic function in multiple thalamic nuclei in *Df(16)1/+* mice,^73^ we predicted that decreased Pf firing underlies the Pf-MSN synaptic transmission deficits. Therefore, we investigated the firing properties of Pf neurons in both genotypes. Surprisingly, we found no differences in the firing or basic membrane properties of Pf neurons between *Df(16)1/+* and WT mice (**Figure S2A–G**), with the exception of higher input resistances in *Df(16)1/+* Pf neurons (**Figure S2H**).

In the absence of decreased firing at the somatic level, our findings suggest a decrease in presynaptic release from Pf terminals onto MSNs. Therefore, we hypothesized that aberrant activation of inhibitory receptors on Pf terminals dampens Pf-MSN synaptic transmission in *Df(16)1/+* mice.^86^ The Pf terminals contain inhibitory M2 muscarinic acetylcholine receptors (mAChR),^87–90^ which are implicated in SCZ symptoms.^91–93^ We therefore tested the role of M2 receptors in Pf-MSN synaptic transmission. Inclusion of the M2 receptor antagonist AF-DX-116 in the ACSF during optogenetic input-output experiments restored *Df(16)1/+* Pf-MSN synaptic transmission and PPR to WT levels (**Figure 3A, B**). Bath application of AF-DX-116 also eliminated the deficit in the frequency of sEPSC events onto MSNs without affecting the amplitude of sEPSC events (**Figure S3A, B**). AF-DX-116 significantly enhanced Pf-MSN synaptic transmission in *Df(16)1/+* mice but reduced it in WT mice (**Figure 3B**), suggesting that M2 receptor-mediated modulation of Pf-MSN synaptic transmission is bidirectional and context-dependent.

**Figure 3.**
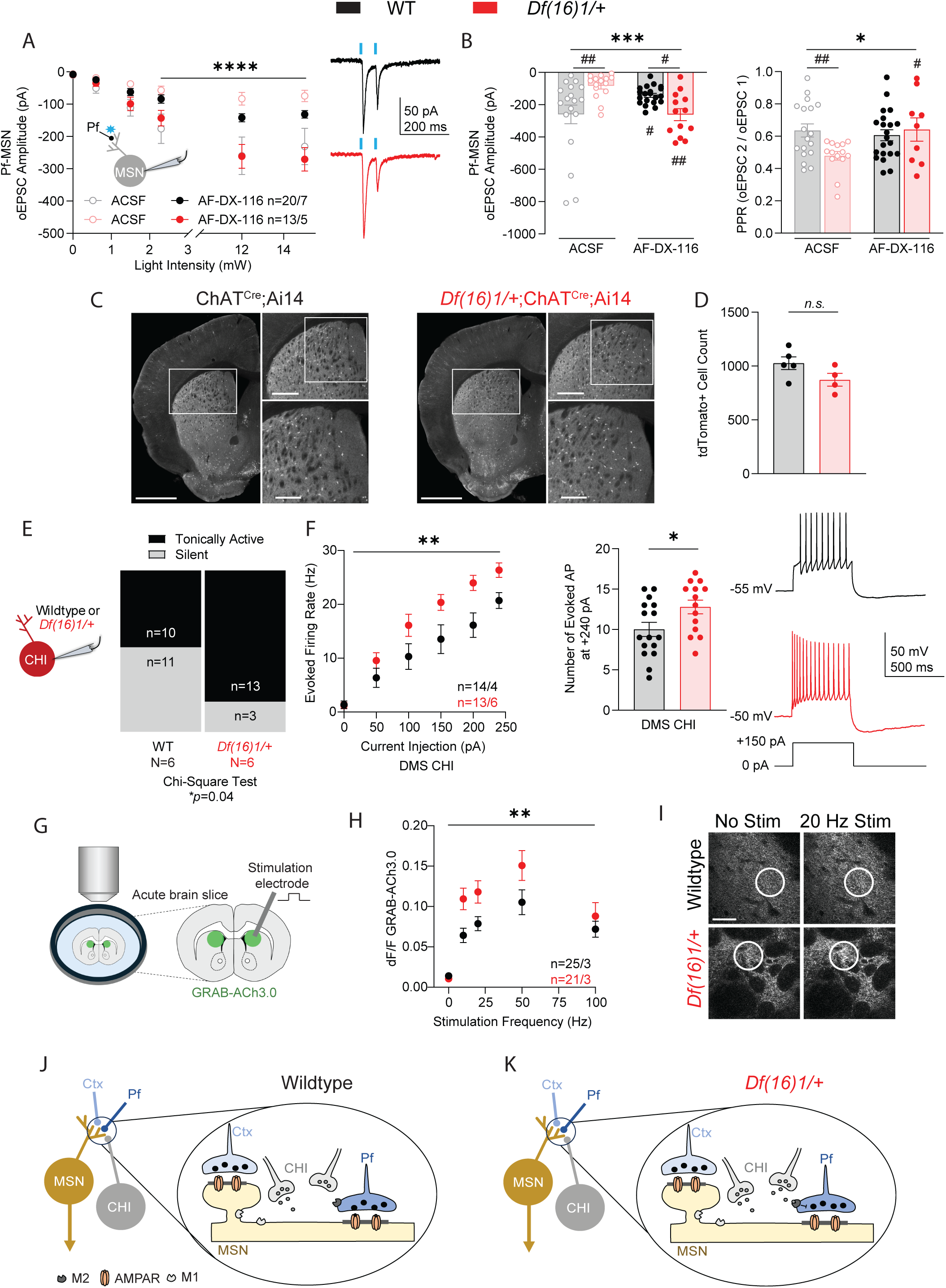
Hypercholinergic state in the DMS of 22q11DS mice. (A) Left: Inclusion of the muscarinic acetylcholine (ACh) M2 receptor antagonist AF-DX-116 in the ACSF (black and red) restored *Df(16)1/+* synaptic transmission at the Pf-MSN synapse compared to control ACSF (gray and pink; mixed-design ANOVA, \*\*\*\**p* < 0.0001). Inset: Schematic of the recording configuration. Right: Example traces of optogenetically evoked Pf-MSN EPSCs in the presence of AF-DX-116. (B) Left: Bath application of AF-DX-116 rescued *Df(16)1/+* Pf-DMS oEPSC amplitude in response to a 12-mW stimulus intensity (two-way ANOVA, \*\*\**p* = 0.0002; Fisher’s LSD test between genotypes, ##*p* = 0.001; between genotypes in the presence of AF-DX-116, #*p* = 0.03; WT AF-DX-116 compared to ACSF, #*p* = 0.02; *Df(16)1/+* AF-DX-116 compared to ACSF, ##*p* = 0.002). Right: Bath application of AF-DX-116 restored *Df(16)1/+* Pf-MSN PPR to WT levels (two-way ANOVA, \**p* = 0.03; Fisher’s LSD test between genotypes, ##*p* = 0.008; between genotypes in the presence of AF-DX-116, *p* = 0.58; WT AF-DX-116 compared to ACSF, *p* = 0.57; *Df(16)1/+* AF-DX-116 compared to ACSF, #*p* = 0.02). (C) Left: Immunofluorescence of tdTomato expression under the choline acetyltransferase (ChAT) promoter in WT mice at lower (left) and higher (right top, bottom) magnifications. Right: Immunofluorescence of ChAT-driven tdTomato in *Df(16)1/+* mice at lower (left) and higher (right top, bottom) magnifications. Scale bars for left images: 1 mm. Scale bars for right top: 0.5 mm. Scale bars for right bottom: 0.25 mm. White boxes show the area of higher magnification. (D) The total number of DMS tdTomato^+^ cholinergic interneurons (CHI) was similar between genotypes (unpaired *t* test, *p* = 0.11). (E) Left: Schematic of the recording configuration. Right: There were more tonically active CHIs in *Df(16)1/+* mice than in WT littermates (Chi-square test, \**p* = 0.04). *n* = number of cells; *N* = number of mice. (F) Left: The evoked firing rate in response to a depolarizing current injection of CHIs from *Df(16)1/+* mice was higher than that from WT littermates (mixed-design ANOVA, \*\**p* = 0.004). Center: The average number of evoked AP in response to a +240-pA current injection was higher in *Df(16)1/+* mice compared to WTs (unpaired *t* test, \**p* = 0.03). Left: Example traces showing the response of CHIs from both genotypes to a depolarizing current injection (+150 pA). (G) Schematic showing the imaging and experimental configuration. (H) There was a greater change in fluorescence (dF/F) of the ACh sensor GRAB-ACh in response to increasing stimulation frequency in *Df(16)1/+* mice compared to WT littermates (mixed-design ANOVA, \*\**p* = 0.009). *n* = number of fields of view or number of mice. (I) One example region of interest from WT and *Df(16)1/+* DMS expressing GRAB-ACh during no stimulation (left) and 20 Hz electrical stimulation (right) is highlighted in the white circle. Scale bar: 35 µm. (J) Schematic representing the Pf-CHI-MSN trisynaptic circuit in WT mice. (K) Schematic representing disruption in synaptic transmission at the Pf-CHI-MSN trisynaptic circuit in *Df(16)1/+* mice. Data shown are mean ± SEM with individual data points overlaid in (B), (D), (F). *n* = the number of cells/number of mice, unless otherwise noted. Blue star (A) represents optical stimulation. Blue lines (A) represent the optical stimulus *n.s.*: not significant. See **Figure S3** for additional electrophysiologic measurements.

M2 modulation of Pf-MSN synaptic transmission may depend on the level of ACh present. The primary source of ACh in the DMS is large, aspiny CHIs.^94,95^ These cells provide a basal tone of ACh within the DMS through tonic firing.^96^ To investigate whether overactivation of M2 mAChRs on Pf terminals in 22q11DS mice arises from aberrant DMS CHI activity, we generated a reporter mouse line that expresses the fluorescent protein tdTomato in CHIs (ChAT^Cre^;Ai14). We then crossed that line with the *Df(16)1/+* mice (*Df(16)1/+*;ChAT^Cre^;Ai14) (**Figure 3C**). We found that, despite similar numbers of CHIs in the DMS (**Figure 3D**), there were significantly more tonically active CHIs in slices from *Df(16)1/+* mice compared to WT littermates (**Figure 3E**). Moreover, when driven to fire through the injection of depolarizing current, CHIs from *Df(16)1/+* mice fired at a higher frequency (**Figure 3F**). There were no additional differences in physiologic properties of CHIs between genotypes (**Figure S3C–I**). These findings suggest that striatal cholinergic tone is greater in *Df(16)1/+* mice. To test this, we investigated the downstream effects of enhanced CHI firing by measuring ACh levels with the GPCR-activation-based (GRAB)-ACh3.0 fluorescent sensor^97^ (**Figure 3G**). Electrically evoked ACh levels were enhanced in acute slices derived from *Df(16)1/+* mice compared to WT littermates (**Figure 3H, I**).

We therefore hypothesized that enhanced CHI firing causes the hypercholinergic state and aberrant activation of presynaptic M2 receptors on Pf terminals. This overactivation may cause a decrease in Pf-MSN synaptic transmission, which leads to amotivation in 22q11DS mice (**Figure 3J, K**).

### Enhancing CHI activity produces amotivation

To evaluate the influence of hyperactive CHIs on motivated behavior, we expressed the Cre-dependent excitatory DREADD receptor hM3Dq in DMS CHIs of ChAT^Cre^ mice (**Figure 4A, B**). Compared to vehicle injection, activating CHIs with C21 injection resulted in fewer lever presses and rewards earned on both PR tasks (**Figure 4C–E**). The reduction in lever pressing on the PR+4 task after CHI activation was smaller (one-sample *t* test with µ = –0.455, *p* = 0.008) than that between *Df(16)1/+* and WT mice (**Figure 4D** and **Figure 1D**), similar to our findings in **Figure 2D**.

**Figure 4.**
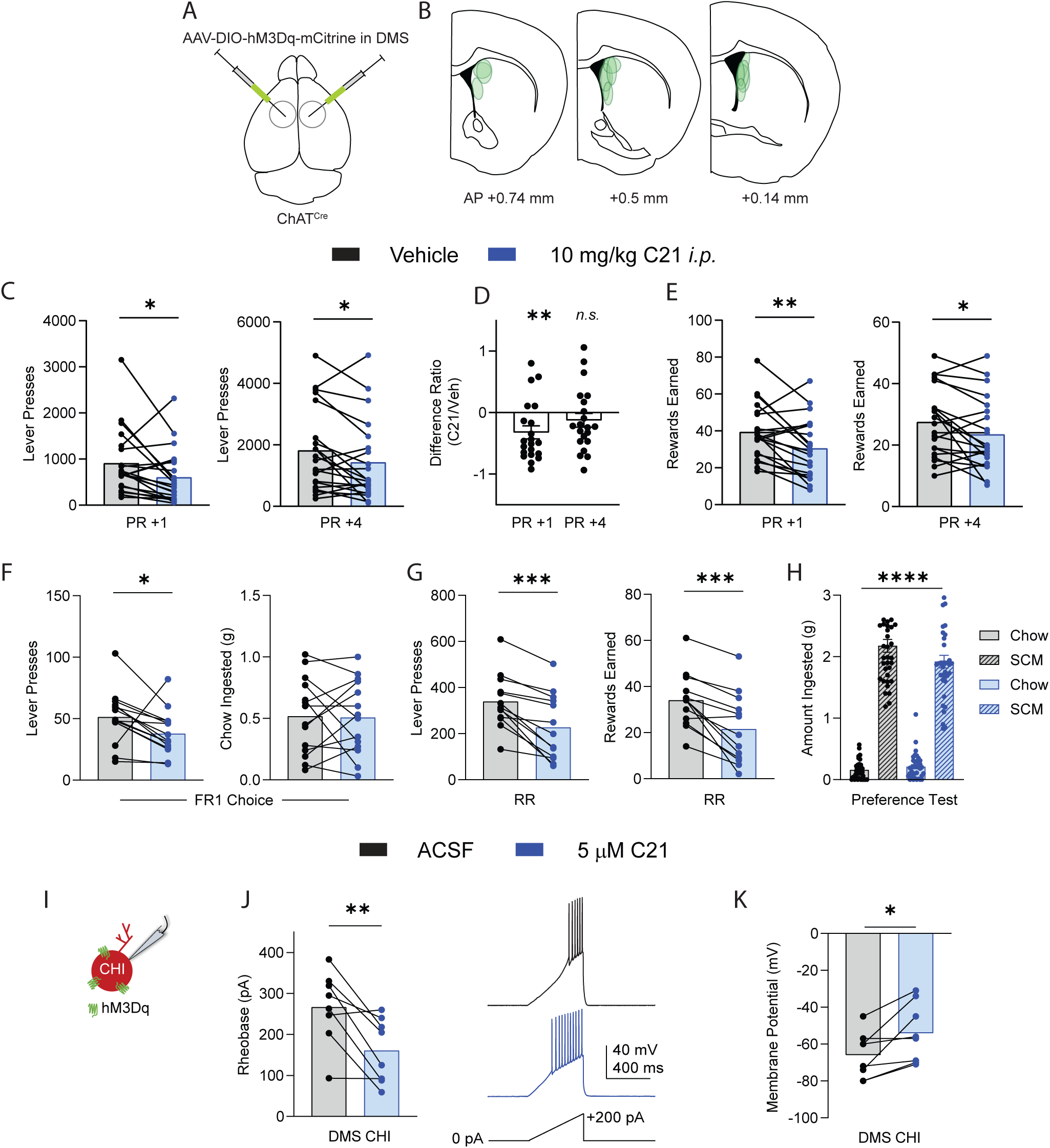
Chemogenetic enhancement of CHI activity produces amotivation. (A) Schematic of the injection paradigm. A Cre-dependent virus containing the excitatory DREADD receptor hM3Dq was bilaterally injected into the DMS of ChAT^Cre^ mice. (B) Histologic verification of injection sites in the DMS (AP +0.74 to +0.14 mm from bregma). Circles are virus expression from individual mice. (C) Mice injected with the DREADDs agonist C21 lever pressed less than vehicle-injected mice in the PR+1 task (left; two-way RM ANOVA,* *p* = 0.04) and in the PR+4 task (right; two-way RM ANOVA, \**p* = 0.03). (D) The average difference ratio between vehicle and C21 injection was –0.32 ± 0.11 in PR+1 task performance and –0.13 ± 0.11 in PR+4 performance. The PR+1 difference ratio was significantly different from a theoretical mean of 0 (one-sample *t* test, \*\**p* = 0.007), but the PR+4 difference ratio was not (one-sample *t* test, *p* = 0.28). (E) The number of rewards earned on the PR+1 (left) and PR+4 (right) tasks was lower after C21 injection than vehicle injection (two-way RM ANOVA for PR+1, \*\**p* = 0.005; for PR+4, \**p* = 0.02). (F) Left: There were significantly fewer lever presses in the FR1 choice task after C21 injection compared to vehicle (two-way RM ANOVA, \**p* = 0.02). Right: There was no difference in the amount of freely available rodent chow ingested in the FR1 choice task between C21 and vehicle injection (two-way RM ANOVA, *p* = 0.89). (G) Both lever presses (left) and the rewards earned (right) on the RR task after C21 injection were significantly different from vehicle injection (two-way RM ANOVA for lever presses, \*\*\**p* = 0.0001; for rewards earned, \*\*\**p* = 0.0001). (H) Mice preferred SCM over standard rodent chow when both were freely available regardless of injection type (three-way ANOVA, \*\*\*\**p* < 0.0001). There was no difference in SCM consumption based on injection type (Tukey’s multiple comparison, *p* = 0.56). (I) Schematic of the recording configuration. Green receptors represent hM3Dq expression. (J) Left: The rheobase current required to induce AP firing in CHIs was lower after application of C21 compared to vehicle (paired *t* test, \*\**p* = 0.008). Right: Example traces of CHIs in the presence of ACSF or C21 responding to a depolarizing current injection ramp, from 0 pA to +200 pA. (K) The membrane potential of CHIs expressing hM3Dq was depolarized after application of C21 compared to vehicle (paired *t* test, \**p* = 0.02). All data shown are mean ± SEM with individual data points overlaid in (C–K). Unless noted, there were no sex differences. *n.s.*: not significant.

Activating CHIs with C21 also altered the performance of mice on the FR and RR tasks (**Figure 4F, G**). In the FR1 choice task, C21 injection resulted in fewer lever presses for SCM but did not change the consumption of freely available chow (**Figure 4F**). Moreover, C21 injection resulted in fewer lever presses and rewards earned on the RR task but did not alter SCM preference (**Figure 4G, H**).

To verify the function of hM3Dq used in the behavioral studies, we made coronal slices containing the DMS of ChAT^Cre^ mice injected with Cre-dependent hM3Dq. We recorded directly from CHIs expressing hM3Dq and measured the change in membrane and firing properties after bath application of C21 (**Figure 4I–K**). As expected, C21 application decreased the rheobase current needed to evoke action potentials (APs) in CHIs (**Figure 4J**) and depolarized the resting membrane potential (**Figure 4K**).

### Pf-MSN synaptic transmission is bidirectionally altered by cholinergic activity in the DMS

We next sought to verify the involvement of CHI hyperactivity in Pf-MSN synaptic transmission. To do this, we recorded Pf-MSN synaptic transmission from ChAT^Cre^ mice with hM3Dq expression in DMS CHIs (**Figure 5A**). Consistent with our hypothesis that hyperactive CHIs weaken Pf-MSN synaptic transmission through M2 receptor activation, bath application of C21 to activate CHIs decreased the Pf drive of MSNs (**Figure 5B, C**). Moreover, consistent with a change in presynaptic release, CHI activation with C21 increased the PPR at the Pf-MSN synapse compared to baseline (**Figure 5D**).

**Figure 5.**
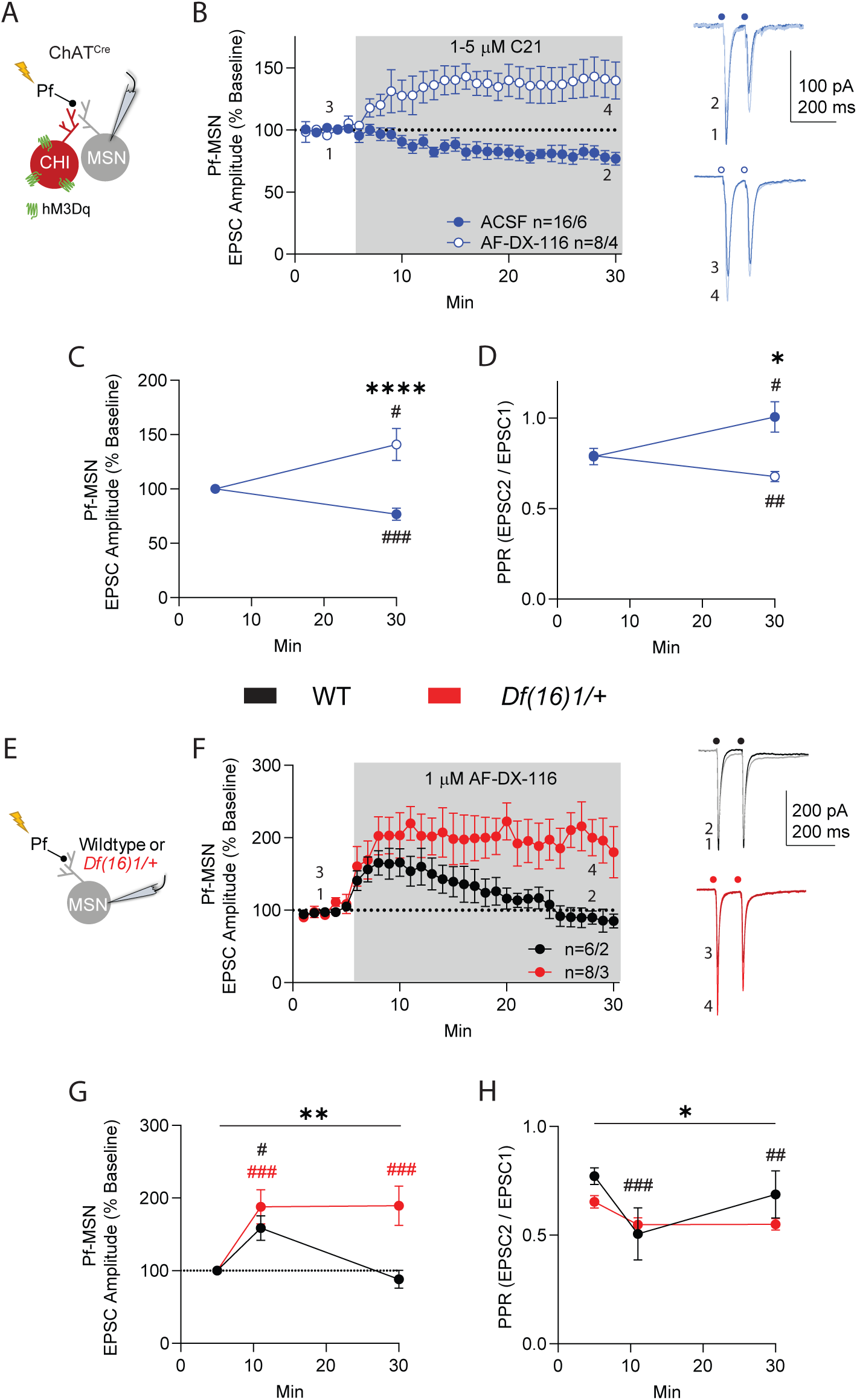
Pf-MSN synaptic transmission is bidirectionally altered by cholinergic transmission. (A) Schematic of the recording configuration. Green receptors represent hM3Dq expression. (B) Left: Activation of DMS CHIs with bath application of C21 decreased Pf-MSN synaptic transmission over time (filled circles). Bath application of C21 in the presence of AF-DX-116 enhanced Pf-MSN synaptic transmission (open circles). Shaded region depicts the bath application of 1-5 µM C21. Right, closed circles: Representative traces demonstrating the decrease in amplitude following C21 application (2) compared to baseline (1). Right, open circles: Representative traces demonstrating an increase in EPSC amplitude following C21 application in the presence of AF-DX-116 relative to baseline (4 and 3, respectively). (C) There was a significant difference in Pf-MSN synaptic transmission in the presence or absence of AF-DX-116 after C21 application (unpaired *t* test of min 25-30, \*\*\*\**p* < 0.0001). The changes in synaptic transmission following C21 application with and without AF-DX-116 present were different from their baseline values (one-sample *t* test with µ = 100 of min 25-30, ###*p* = 0.0009 for ACSF; #*p* = 0.028 for AF-DX-116). (D) There was a significant difference in the PPR between electrically evoked EPSCs at the Pf-MSN synapse after C21 wash in the presence or absence of AF-DX-116 (paired *t* test of min 25-30, #*p* = 0.039 for ACSF; ##*p* = 0.004 for AF-DX-116). There was a significant difference in the change in PPR at min 25-30 between groups (data analyzed by unpaired *t* test, \**p* = 0.01). (E) Schematic of the recording configuration. (F) Left: Bath application of AF-DX-116 temporarily enhanced Pf-MSN synaptic transmission in WT mice. In *Df(16)1/+* mice, bath application of AF-DX-116 led to a prolonged increase in Pf-MSN synaptic transmission. Shaded region depicts bath application of 1 µM AF-DX-116. Right: Representative traces showing no change in WT or the increase in *Df(16)1/+* mice in Pf-MSN synaptic transmission after AF-DX-116 application. (G) There was a significant difference in the response to AF-DX-116 application based on genotype and time following application (two-way RM ANOVA for min 1-5 (5), 6-11 (11), and 25-30 (30), *\*\*p* = 0.009; Tukey’s multiple comparison test min 5 vs min 11 for wildtype, *#p* = 0.046; for *Df(16)1*/+ *###p* = 0.0006. For *Df(16)1/+* min 5 vs min 30 ###*p* = 0.0005). (H) There was a significant difference in the PPR in response to AF-DX-116 application based on genotype and time following application (two-way RM ANOVA, \**p* = 0.034; Tukey’s multiple comparison test min 5 vs 11 for wildtype, ###*p* = 0.0001; for min 11 vs 30 for wildtype ##*p* = 0.006). Data shown are mean ± SEM. *n* = the number of cells/number of mice. Lightning bolts (A, E) represent electrical stimulation. Circles (B, F), right, represent stimulus artifacts.

Interestingly, the M2 receptor antagonist AF-DX-116 not only occluded the inhibitory effect of C21 on Pf-MSN synaptic transmission but even reversed it. In the presence of AF-DX-116, C21 enhanced Pf-MSN synaptic transmission (**Figure 5B, C**) and led to a significant decrease in Pf-MSN PPR (**Figure 5D**). These findings show that CHI activity bidirectionally affects Pf-MSN synaptic transmission, and this effect depends on M2 receptor availability.

Given that CHIs are tonically more active and release more ACh in the DMS in 22q11DS mice compared to WT mice, we hypothesized that relieving the inhibitory hypercholinergic effect on Pf-MSN synaptic transmission will strengthen these synapses, and this strengthening will be more pronounced in 22q11DS mice than in WT mice. Indeed, prolonged (25-minute) application of AF-DX-116 strengthened Pf-MSN synaptic transmission in oblique horizontal slices (**Figure 5E-G**). This strengthening was substantially stronger in 22q11DS mice compared to WT littermates. In WT mice, the evoked EPSC amplitude returned to baseline levels by minutes 25–30 (**Figure 5F, G**). However, in *Df(16)1/+* mice, AF-DX-116 application led to a stronger, sustained enhancement of Pf drive of MSNs (**Figure 5F, G**). There was a significant interaction between genotype and time following AF-DX-116 application for PPR at the Pf-MSN synapse, suggesting the enhancement of synaptic transmission at this synapse is presynaptically mediated (**Figure 5H**).

### Gene expression is altered in CHIs from the dorsal striatum of *Df(16)1/+* mice

After verifying that CHI activity influences Pf-MSN synaptic transmission, we investigated gene expression changes in *Df(16)1/+* CHIs that might contribute to their hyperactivity. To this end, we performed snRNA-seq on the dorsal striatum of four *Df(16)1/+* and four WT littermates (**Figure 6A**). Cell types were identified as outlined in **Figure S4A**. After 43,319 nuclei passed quality control (**Figure S4B**), VoxHunt mapping onto *Allen Developing Mouse Brain Atlas* data revealed that 4 clusters were derived from cortex or thalamus (**Figure S4C**), likely due to contamination during dissection of the dorsal striatum. These clusters were removed from further analysis, leaving 35,167 nuclei derived from dorsal striatum (**Figure 6B**). We then identified glial cell types (**Figure S4D**) using previously reported marker genes (**Table S1**). Of the remaining neuronal nuclei, 86% were derived from MSNs expressing the MSN markers *Rgs9* and *Ppp1r1b* (**Figure S4E**) and were split into 5 clusters: matrix D1-dopamine receptor containing, matrix D2-dopamine receptor containing, patch D1, patch D2, and eccentric MSNs.

**Figure 6.**
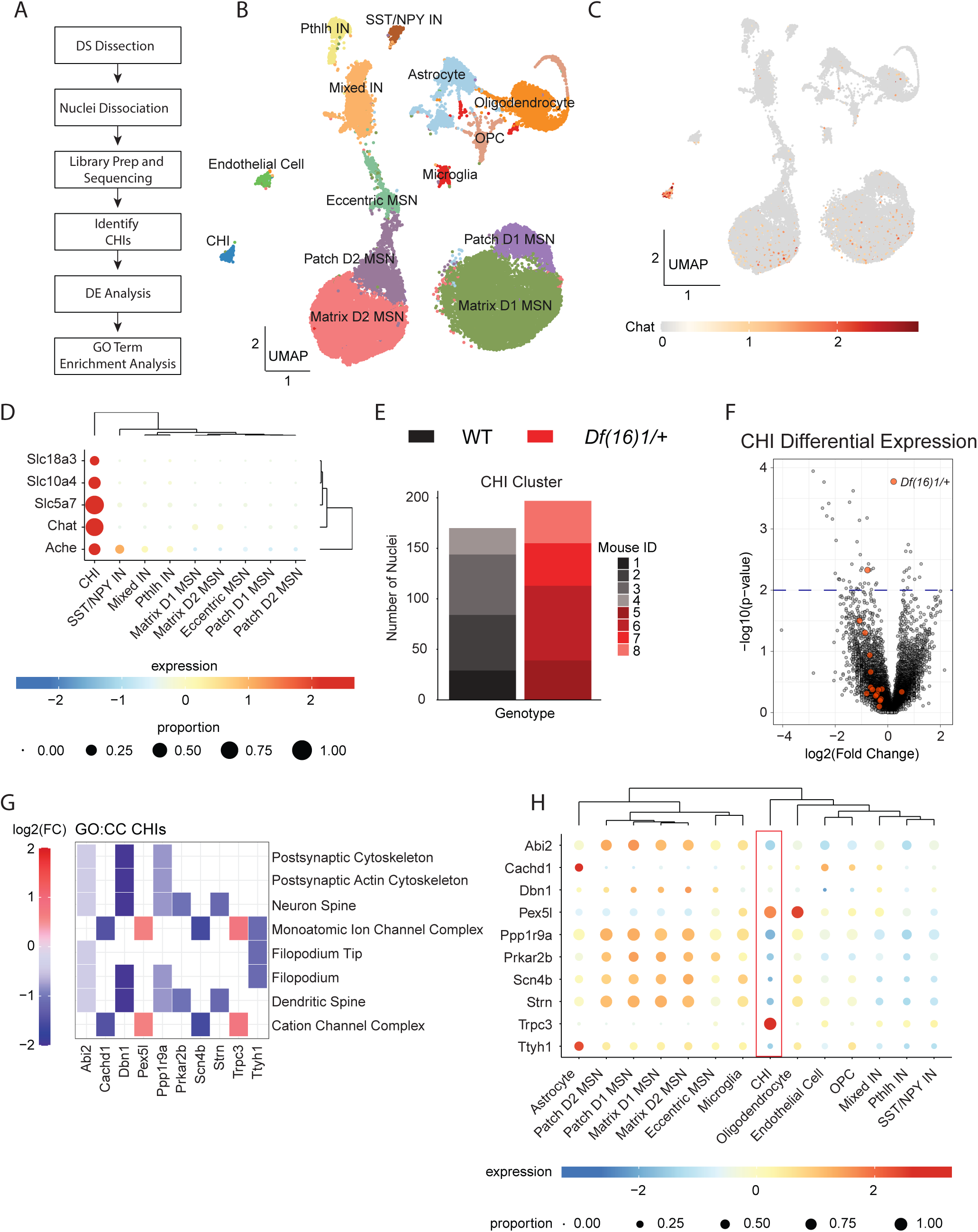
Gene expression is altered in CHIs from the dorsal striatum of 22q11DS mice. (A) Overview of single-nucleus RNA-sequencing (snRNA-seq) analysis. (B) Uniform Manifold Approximation and Projection (UMAP) plot with cluster annotations indicated by color. (C) UMAP plot of the CHI marker, *Chat*. Color indicates the normalized transcript level. (D) Dot plot showing CHI marker expression among neuronal clusters. (E) Bar plot of the number of CHI nuclei detected per mouse. Color indicates mouse identity. (F) Volcano plot showing differential gene expression in *Df(16)1/+* CHIs compared to WT littermates. Red circles indicate genes within the *Df(16)1/+* hemizygous region. (G) Heatmap showing cellular compartment (CC) terms overrepresented among transcripts differentially expressed in *Df(16)1/+* CHIs. Color indicates log2(Fold Change) of each transcript. (H) Dot plot showing normalized expression of each transcript from (G) within each cell type found in our dorsal striatal data. Red box highlights the CHI cluster. expression: normalized average expression; proportion: proportion of cells expressing a marker within a cluster. Data in (A–H) were produced by snRNA-seq analysis of 35,167 nuclei derived from the dorsal striatum of 4 *Df(16)1/+* mice and 4 WT littermates. Nuclei derived from cortex and thalamus were removed before analysis. See **Figure S4** for additional snRNA-seq analyses and **Tables S1 and S2** for supporting data.

These clusters were characterized based on previously reported markers of each group (**Figure S4F).**^98–101^ The remaining nuclei primarily consisted of GABAergic interneurons expressing *Pvalb* (**Figure S4G**), *Cck* (**Figure S4H**), *Pthlh* (**Figure S4I**), or co-expressing *Sst* and *Npy* (**Figure S4J**). Finally, a small cluster of cells expressing *Chat* and other genes involved in ACh degradation (*Ache*), transport (*Slc18a3* and *Slc10a4*), or synthesis (*Slc5a7*) were identified as CHIs (**Figure 6B–D**). Consistent with previous reports,^102–104^ CHIs represented ∼1% of cells in the dorsal striatum (373 nuclei across both genotypes). The number of CHI nuclei was similar between genotypes, consistent with our immunofluorescence data (**Figure 6E**).

We performed differential gene expression analysis to compare *Df(16)1/+* and WT CHIs. When nuclei were pooled across all clusters, genes in the *Df(16)1/+* hemizygous region were downregulated in *Df(16)1/+* nuclei, as expected (**Figure S4K**). Similarly, most genes in the *Df(16)/1+* hemizygous region were downregulated in *Df(16)1/+* CHIs (**Figure 6F**), but only *Tango2* met the significance threshold of *p* < 0.01 (**Table S2**). Gene ontology (GO) enrichment analysis revealed that cellular compartment terms associated with postsynaptic compartments and dendritic spines were overrepresented among downregulated genes (defined as log2(Fold Change) < 0 and *p* < 0.01) (**Figure 6G**). The terms “monoatomic ion channel complex” and “cation channel complex” were also overrepresented among differentially expressed genes (defined as *p* < 0.01). Both terms include two genes (*Pex5l,* which encodes TRIP8b, an auxiliary subunit of hyperpolarization-activated cyclic nucleotide-gated [HCN] channels and *Trpc3*, which encodes the non-selective Ca^2+^-permeable transient receptor potential 3 channel [TRPC3]) that were increased in *Df(16)1/+* CHIs and highly expressed in CHIs relative to other cell types in the dorsal striatum (**Figures 6G, H**). Finally, we found that the cellular composition of the dorsal striatum was similar between genotypes (**Figure S4L, M**). Therefore, CHIs from *Df(16)1/+* mice exhibit altered expression of transcripts encoding ion channels that may contribute to their hyperactive state.

### Manipulating CHI activity and Pf-DMS synaptic transmission in 22q11DS mice is not sufficient to rescue amotivation

Finally, we tested the hypothesis that restoring CHI activity and Pf-DMS synaptic transmission in 22q11DS mice to WT levels, either separately or jointly, will rescue the motivational impairment. We first decreased abnormally high CHI activity in 22q11DS mice by expressing hM4Di in DMS CHIs in *Df(16)1/+*;ChAT^Cre^ mice (**Figure 7A, B**). However, C21 injection to decrease CHI activity did not increase lever pressing in these mice on the PR+1 or PR+4 tasks (**Figure 7C–E**). Similarly, C21 injection did not change performance on the RR task or alter the preference for SCM over chow (**Figure S5A, C**), but did decrease lever presses in the FR1 task (**Figure S5B**).

**Figure 7.**
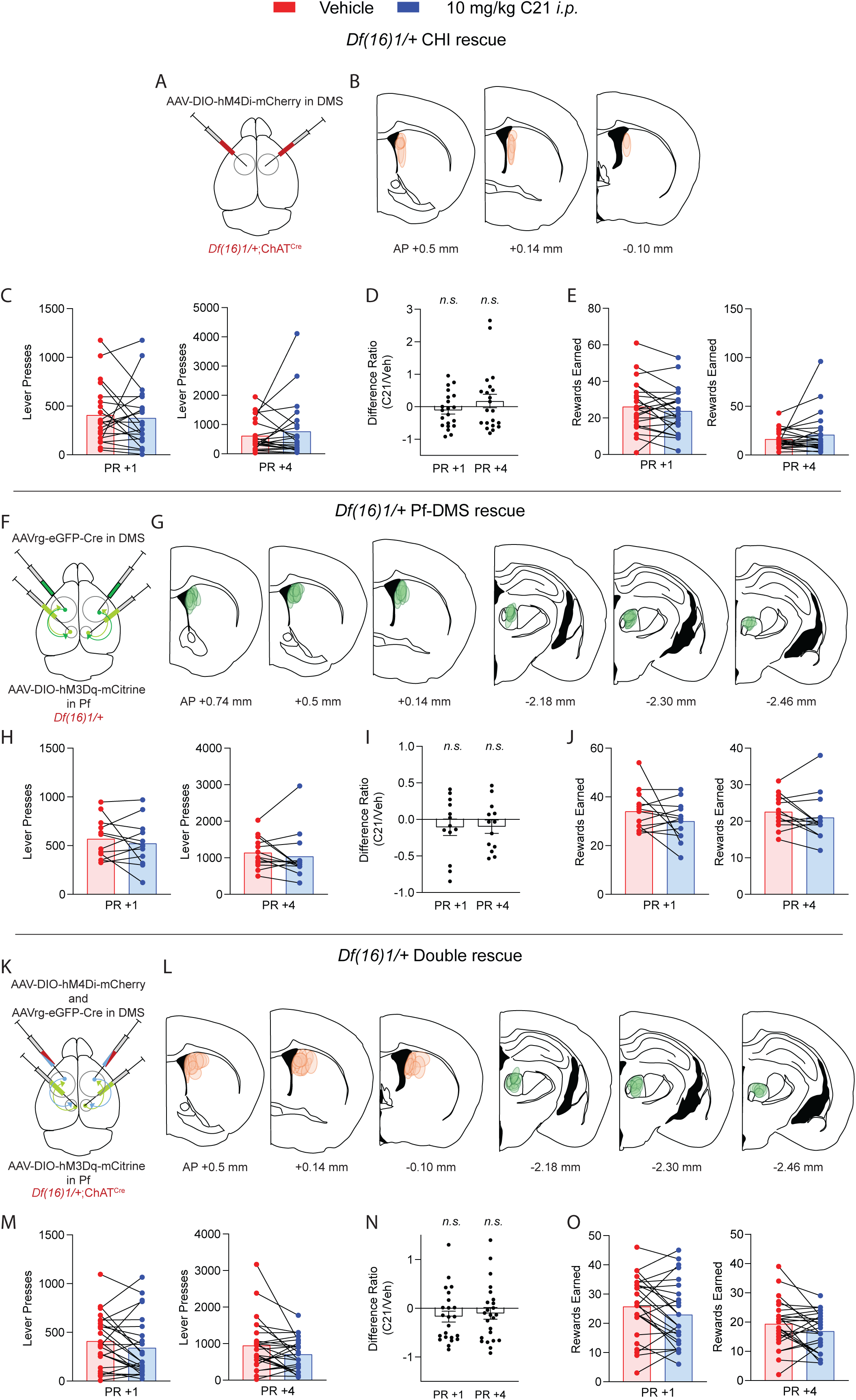
Manipulating CHI activity and Pf-DMS synaptic transmission in 22q11DS mice is not sufficient to rescue amotivation. (A) Schematic of the injection paradigm. A Cre-dependent virus containing the inhibitory DREADD receptor hM4Di was bilaterally injected into the DMS of *Df(16)1/+*;ChAT^Cre^ mice. (B) Histologic verification of injection sites into the DMS (AP +0.5 to –0.10 mm from bregma). Circles are virus expression from individual mice. (C) Injection of C21 did not affect performance on PR+1 (left) or PR+4 (right) in *Df(16)1/+* mice (two-way RM ANOVA for PR+1, *p* = 0.98; for PR+4, *p* = 0.45). (D) The difference ratio in the PR+1 and PR+4 task performance between vehicle and C21 injection was not significantly different from 0 (one-sample *t* test for PR+1, –0.11 ± 0.12, *p* = 0.37; for PR+4, 0.18 ± 0.21, *p* = 0.41). (E) The number of rewards earned on the PR+1 (left) and PR+4 (right) tasks were not significantly different between vehicle and C21 injection (two-way RM ANOVA for PR+1, *p* = 0.38; for PR+4, *p* = 0.21). (F) Schematic of the injection paradigm. A retrogradely-transported virus containing Cre was bilaterally injected into the DMS of *Df(16)1/+* mice. A Cre-dependent virus containing the excitatory DREADD receptor hM3Dq was bilaterally injected into the Pf of the same mouse. (G) Histologic verification of the injection sites in the DMS (AP +0.74 to +0.14 mm) and Pf (AP –2.18 to –4.26 mm). Circles are virus expression from individual mice. (H) Injection of C21 did not alter performance on the PR+1 (left) or PR+4 (right) tasks in *Df(16)1/+* mice (two-way RM ANOVA for PR+1, *p* = 0.49; for PR+4, *p* = 0.49). (I) The difference ratio in the PR+1 and PR+4 task performance between vehicle and C21 injection is not different from 0 (one-sample *t* test for PR+1, –0.11 ± 0.11, *p* = 0.36; for PR+4, – 0.1 ± 0.09, *p* = 0.3). (J) There was no difference in the number of rewards earned for PR+1 (left) or PR+4 (right) in *Df(16)1/+* mice after C21 injection compared to vehicle (two-way RM ANOVA for PR+1, *p* = 0.23; for PR+4, *p* = 0.24). (K) Schematic of the injection paradigm. A retrogradely-transported Cre virus and a virus containing the Cre-dependent inhibitory DREADD receptor hM4Di were bilaterally injected into the DMS of *Df(16)1/+*;ChAT^Cre^ mice. In the same mice, a virus containing a Cre-dependent excitatory DREADD receptor hM3Dq was bilaterally injected into the Pf. (L) Histologic verification of the injection sites in the DMS (AP +0.5 to –0.1 mm) and Pf (AP – 2.18 to –2.46 mm). Circles are virus expression from individual mice. (M) Injection of C21 did not affect *Df(16)1/+* mouse performance on the PR+1 task (left; two-way RM ANOVA, *p* = 0.08) or the PR+4 (right; two-way RM ANOVA, *p* = 0.08). (N) The difference ratio for PR+1 and PR+4 task performance between vehicle and C21 injection was not different from 0 (one-sample *t* test for PR+1, –0.17 ± 0.11, *p* = 0.14; for PR+4, –0.1 ± 0.12, *p* = 0.39). (O) C21 injection did not affect the number of rewards earned during the PR+1 or PR+4 tasks (left and right, respectively; two-way RM ANOVA for PR+1, *p* = 0.08; for PR+4, *p* = 0.18). Data shown are mean ± SEM with individual data points overlaid in (C–E), (H–J), (M–O). Unless noted, there were no sex differences. *n.s.*: not significant. See **Figure S5** for additional behavioral measurements.

Next, we attempted to rescue Pf-DMS synaptic transmission in *Df(16)1/+* mice by expressing hM3Dq in the Pf-DMS pathway (**Figure 7F, G**). C21 injection to increase Pf-DMS synaptic transmission did not rescue amotivation in either PR task in these mice (**Figure 7H–J**). C21 injection neither altered performance on the RR task (**Figure S5D**) nor changed the willingness to work for SCM over food chow in the FR1 choice task (**Figure S5E, F**). Similarly, C21 injection did not change the preference for SCM over food chow (**Figure S5G**).

Finally, in the “double rescue” experiments, dual expression of hM4Di in DMS CHIs and hM3Dq in the Pf-DMS pathway in *Df(16)1/+*;ChAT^Cre^ mice (**Figure 7K, L**) failed to rescue amotivation in the PR tasks (**Figure 7M–O**). C21 injection did not change RR or FR1 choice task performance (**Figure S5H–J**) or SCM preference (**Figure S5K**), suggesting the Pf-DMS pathway, while dysregulated in *Df(16)1/+* mice, is not solely responsible for the amotivation state.

## Discussion

Here, using a mouse model of 22q11DS, we implicate the Pf-DMS circuit and its modulation by the striatal CHIs in motivational states. To our knowledge, this is the first time that this circuit has been linked to motivated behaviors. Our data argue that the hypercholinergic state triggered by CHI hyperactivity overly inhibits glutamatergic synaptic transmission at Pf-DMS synapses and that this mechanism is at least partially responsible for amotivation in 22q11DS mice. Therefore, our findings broaden the understanding of motivation processing in native and pathologic states such as 22q11DS.

Thalamostriatal signaling is known to participate in attentional processing of incoming sensory information.^50,105,106^ The intralaminar thalamic nuclei, which contain the Pf, encode salient sensory information through a burst of activity onto striatal CHIs.^107^ In response, CHIs burst fire and release ACh, which activates nicotinic AChRs on passing dopaminergic boutons to release dopamine. Dopamine acts on inhibitory D2 receptors on CHIs to pause their firing. This pattern of CHI activity is known as a burst-pause pattern of spiking and is believed to function as a saliency signal.^107^ Our findings deviate from this traditional view by implicating the Pf-MSN-CHI trisynaptic circuit in motivated behaviors. Moreover, unlike previous findings that the Pf influences CHI activity, our data show that CHI activity affects the Pf drive of MSNs through the activation of M2 AChRs. We show that in a hypercholinergic state in 22q11DS or induced by excitatory DREADDs, blocking the M2 receptor causes a prolonged enhancement of Pf-MSN synaptic transmission. Whether this enhanced Pf-MSN synaptic transmission would rescue amotivation in 22q11DS mice remains unknown.

Our snRNA-seq analysis in CHIs from *Df(16)1/+* mice revealed differential expression of multiple genes involved in regulating membrane depolarization including notably higher expression of *Trpc3* and *Pex5l* in *Df(16)1/+* CHIs than in controls. *Trpc3* encodes TRPC3, a non-selective Ca^2+^-permeable channel that is expressed in striatal CHIs and is the target of metabotropic glutamate receptor signaling^108^ and dopamine D5-receptor signaling.^109^ These channels are G-protein coupled receptor–Ca^2+^ signaling effectors in neurons.^110^ Importantly, TRPC3 has been shown to modulate the depolarization and tonic firing of CHIs, as TRPC3 blockade reduces the firing rate of striatal CHIs.^109^ Thus, the overexpression of TRPC3 in 22q11DS CHIs could cause increased tonic firing and increased firing rates, which is consistent with our findings. *Pex5l* encodes TRIP8b, an auxiliary subunit of HCN channels. HCN channels are permeable to Na^+^ and K^+^ and mediate I_h_, which influences resting membrane potential and input resistance.^111^ As the reversal potential of HCN channels is between –50 and –20 mV,^112^ activation of these channels at those membrane potentials depolarizes the neuron, positioning it closer to the AP threshold. Moreover, TRIP8b regulates the subcellular distribution of HCN channels.^113,114^ Overexpression of TRIP8b in CHIs could promote the clustering of HCN channels to alter the integration of incoming excitatory synaptic transmission, further promoting CHIs to be in the active state. The elevated expression level of either or both genes may explain the increase in tonic CHI activity in 22q11DS mice.

Mimicking the altered DMS physiology (i.e., with chemogenetic Pf-DMS disruption or chemogenetic CHI hyperactivation) observed in 22q11DS mice produces amotivation in WT mice, suggesting that the Pf-DMS projections and CHI-mediated ACh release that modulates synaptic transmission in Pf-DMS projections contribute to motivation control. The mimicking methods used here caused amotivation on a smaller scale than what occurs in 22q11DS. Thus, 22q11DS likely disrupts multiple neural circuits involved in appetitive food reward behavior. This is consistent with our inability to rescue the motivational deficits in 22q11DS mice by normalizing Pf-DMS synaptic transmission, CHI firing, or both. These negative results exemplify the complexity of motivation as an underlying behavioral process. Indeed, multiple neural circuits are implicated in motivated behavior. Dopamine signaling within the nucleus accumbens is necessary for motivational processes.^23,80^ Although we did not observe changes in accumbal physiology, disruptions in Pf-DMS signaling may decrease dopamine release within the DMS.^58^ Another example involves the cortical control of motivation, with prefrontal cortices and the hippocampus being integral in associative learning between environmental stimuli and motivated actions.^115–117^ Additionally, the paraventricular nucleus of the thalamus (PVT) is critical mediator of top-down control of cue-motivated behavior in rats.^118^ Through two distinct cellular projections, the PVT innervates the NAc and encodes the execution and termination of motivated goal-directed actions.^119^ Finally, similar to the NAc, the amygdala receives dopaminergic inputs and is involved in reinforcement learning, likely driving motivated behaviors.^120,121^ Deficits within these circuits could additionally contribute to amotivation in 22q11DS, and a full rescue of motivational processes will not occur until all underlying components are restored. However, the role of these mechanisms in 22q11DS amotivation is still unknown.

Nonetheless, we show here that the Pf-DMS circuit and DMS CHIs are necessary for motivated behavior, as the disruption of their normal functioning disrupts motivational processes. Given that CHI activity through ACh release controls Pf-DMS synaptic transmission, the simplest explanation of our results is that hyperactive CHIs in 22q11DS produce a hypercholinergic state in the DMS that inhibits glutamate release at Pf-DMS synapses through overactivation of the M2 AChRs. Our findings implicating M2 mAChRs in amotivation align with decades of preclinical and clinical research on the relationship between mAChRs and SCZ symptoms.^91–93^ Tandon et al.^91^ showed that nonselective mAChR antagonists alleviated negative SCZ symptoms in patients. Moreover, the M1/M4 agonist xanomeline can treat positive and negative symptoms of SCZ.^92^ However, this drug was pulled from clinical trials due to significant gastrointestinal side effects. More recently, the combination of xanomeline and the peripheral mAChR antagonist trospium chloride (KarXT) has shown promising clinical results without the negative side effects.^93^ KarXT effectively ameliorates negative and positive SCZ symptoms in a dopamine-independent manner.^93,122^ Although additional research is needed to identify the cell type and circuit-specific mechanisms underlying the effectiveness of KarXT, our data implicate the Pf-DMS neural circuit as a potential target of this drug to improve negative symptoms.

## Resource Availability

### Lead Contact

Further information and requests for resources and reagents should be directed to and will be fulfilled by the lead contact, Stanislav Zakharenko.

### Materials Availability

Any materials generated in this study are available from the lead contact with a completed materials transfer agreement.

### Data and Code Availability

- The snRNA-seq data have been deposited in GEO as GEO: GSE293229 and are publicly available as of the date of publication.
- The code used for the analysis of snRNA-seq data is available at Github: https://github.com/ZakharenkoLab/Df1_dorsalstriatum_snRNAseq.
- Any additional information required to reanalyze the data reported in this paper is available from the lead contact upon request.

## Acknowledgements

The authors thank Abbas Shirinifard for assistance with StarDist segmentation, Rebekah Doerfler for manuscript editing, and Zakharenko lab members for constructive comments and technical assistance. This work was funded, in part, by National Institutes of Health grants R01MH136215, R01MH139498 (S.S.Z.), K99MH129617 (M.H.P.), the Stanford Maternal and Child Health Research Institute Uytengsu-Hamilton 22q11 Neuropsychiatry Research Program grants UH22QEXTFY21 and UH22QEXTFY23 (S.S.Z.), BBRF Distinguished Investigator Grant from the Brain & Behavior Research Foundation (S.S.Z), and the American Lebanese Syrian Associated Charities (S.S.Z). The content is solely the responsibility of the authors and does not necessarily represent the official views of the National Institutes of Health or other granting agencies.

## Author Contributions

Conceptualization: M.H.P. and S.S.Z.

Electrophysiology, two-photon imaging: M.H.P.

Behavior: B.J.W.T.

Immunofluorescence, histology: M.H.P and A.B.S. snRNA-seq

sample collection: M.H.P. and K.T.T.

snRNA-seq nuclei dissociation and library preparation: A.J.T.

snRNA-seq data processing and analysis: K.T.T., S.L.F., and C.A.R.

Visualization: M.H.P.

Funding acquisition: M.H.P. and S.S.Z.

Supervision: S.S.Z.

Writing – original draft: M.H.P.

Writing – review and editing: S.S.Z.

## Supplemental Figures

**Figure S1. Additional behavioral and electrophysiologic measures in WT and 22q11DS mice (Related to Figure 1).**

(A) *Df(16)1/+* mice earned fewer rewards on the progressive ratio (PR)+1 task (left) and the PR+4 task (right) compared to WT littermates (two-way ANOVA for PR+1, \**p* = 0.01; for PR+4, \*\*\**p* = 0.007).

(B) There was no difference in performance (left) or number of rewards earned (right) between genotypes on the random ratio (RR) task (two-way ANOVA for lever presses, *p* = 0.06; for rewards earned, *p* = 0.09). There was a sex difference in RR performance (two-way ANOVA, \**p* = 0.04).

(C) There was no difference in the amount of freely available rodent chow consumed in the fixed ratio 1 (FR1) choice task after injection of C21 (two-way ANOVA, *p* = 0.69).

(D) There was no difference in the cumulative distance traveled over a 30 min period in an open field between genotypes (mixed-design ANOVA, *p* = 0.73). Inset: There was no difference in the overall distance travelled between genotypes (two-way ANOVA, *p* = 0.18). The animal’s sex affected overall distance travelled (two-way ANOVA, \**p* = 0.046).

(E) There was no difference in the cumulative time spent in the center of the open field arena over a 30 min period between genotypes (mixed-design ANOVA, *p* > 0.99). Inset: There was no difference in the overall time spent in the center of the open field arena (two-way ANOVA, *p* = 0.86).

(F) Left: Schematic of the recording configuration. Center: Electrically evoked excitatory postsynaptic current (EPSC) amplitudes in NAc MSNs in response to increasing stimulus intensity were similar between genotypes (mixed-design ANOVA, *p* > 0.99). Right: Example traces of electrically evoked EPSCs in nucleus accumbens (NAc) medium spiny neurons (MSNs).

(G) Schematic of the recording configuration.

(H) The evoked firing rate in response to increasing current step intensities was similar between genotypes in dorsomedial striatal (DMS) MSNs (mixed-design ANOVA, *p* = 0.08).

(I) There was no difference in the number of evoked action potentials (AP) in response to +60 pA current injection between genotypes in DMS MSNs (unpaired *t* test, *p* = 0.35).

(J) The minimum current injected to reach the AP threshold, or rheobase, was not different between genotypes in DMS MSNs (unpaired *t* test, *p* = 0.63).

(K) Left: Example traces showing the response of DMS MSNs from both genotypes to a depolarizing current injection (+60 pA). Right: Example traces of DMS MSNs from both genotypes responding to a depolarizing current injection ramp, from 0 pA to +200 pA.

(L) The resting membrane potential of DMS MSNs was not different between genotypes (unpaired *t* test, *p* = 0.98).

(M) The AP amplitude of DMS MSNs in response to depolarizing current injection was not different between genotypes (unpaired *t* test, *p* = 0.69).

(N) The AP halfwidth of DMS MSNs was not different between genotypes (unpaired *t* test, *p* = 0.35).

(O) The membrane capacitance of DMS MSNs was not different between genotypes (unpaired *t* test, *p* = 0.93).

(P) The input resistance of DMS MSNs was not different between genotypes (unpaired *t* test, *p* = 0.64).

All data shown are mean ± SEM with individual data points overlaid in (A–C), (D, inset), (E, inset), (I–J), (L–P).

Unless noted, there were no sex differences.

*N* = number of mice. *n* = the number of cells/number of mice.

Lightning bolt (F, left) represents electrical stimulation. Circles (F, right) represent stimulus artifacts.

**Figure S2. Electrophysiologic properties of parafascicular nucleus (Pf) neurons in WT and 22q11DS mice (Related to Figure 2).**

(A) Left: Schematic of the experimental configuration. Right: The evoked firing rate of Pf neurons in response to increasing current step intensities was similar between genotypes (mixed-design ANOVA, *p* = 0.93).

(B) There was no difference in the number of evoked APs in response to +10 pA current injection in Pf neurons between genotypes (unpaired *t* test, *p* = 0.9).

(C) Left: The rheobase in Pf neurons was not different between genotypes (unpaired *t* test, *p* = 0.55). Center: Example traces showing the response of Pf neurons from both genotypes to a depolarizing current injection (+10 pA). Right: Example traces of Pf neurons from both genotypes responding to a depolarizing current injection ramp, from 0 pA to +200 pA.

(D) The resting membrane potential of Pf neurons was not different between genotypes (unpaired *t* test, *p* = 0.74).

(E) The AP amplitude of Pf neurons in response to depolarizing current injection was not different between genotypes (unpaired *t* test, *p* = 0.54).

(F) The AP halfwidth of Pf neurons was not different between genotypes (unpaired *t* test, *p* = 0.4).

(G) The membrane capacitance of Pf neurons was not different between genotypes (unpaired *t* test, *p* = 0.07).

(H) The input resistance of Pf neurons from *Df(16)1/+* was higher than in WT littermates (unpaired *t* test, \**p* = 0.03).

Data shown are mean ± SEM with individual data points overlaid in (B–H).

*n* = the number of cells/number of mice.

**Figure S3. Electrophysiologic properties of DMS cholinergic interneurons (CHIs) in WT and 22q11DS mice (Related to Figure 3).**

(A) Schematic of the recording configuration.

(B) Left: Bath application of the M2 antagonist AF-DX-116 did not affect the amplitude of sEPSCs in DMS MSNs between genotypes (unpaired *t* test, *p* = 0.1). Center: Bath application of AF-DX-116 restored the interevent interval of sEPSCs in *Df(16)1/+* DMS MSNs to WT levels (unpaired *t* test, *p* = 0.09). Right: Example traces of sEPSCs from WT and *Df(16)1/+* MSNs in the presence of AF-DX-116.

(C) Schematic of the recording configuration.

(D) There was no difference in the membrane potential of DMS CHIs between genotypes (unpaired *t* test, *p* = 0.4).

(E) There was no difference in the rheobase of CHIs between genotypes (unpaired *t* test, *p* = 0.93).

(F) There was no difference in the amplitude of evoked APs in response to a depolarizing current injection of CHIs between genotypes (unpaired *t* test, *p* = 0.15).

(G) There was no difference in the AP halfwidth of CHIs between genotypes (unpaired *t* test, *p* = 0.63).

(H) There was no difference in the CHI membrane capacitance between genotypes (unpaired *t* test, *p* = 0.74).

(I) The input resistance of CHIs was not different between genotypes (unpaired *t* test, *p* = 0.11).

Data shown are mean ± SEM with individual data points overlaid in (B), (D–I).

**Figure S4. Single-nucleus RNA-sequencing (snRNA-seq) analysis of the dorsal striatum of 22q11DS mice (Related to Figure 6).**

(A) Right: Overview of the workflow for cell type identification in snRNA-seq data, with box color indicating the figure (left) containing relevant data.

(B) UMAP plot with cluster annotation indicated by color. Plot contains nuclei derived from the thalamus and cortex, which were unintended contaminants from dissection of the dorsal striatum. These clusters were removed prior to downstream analyses.

(C) VoxHunt correlation analysis mapping selected snRNA-seq clusters from (B) onto the P56 mouse brain. The schematic indicates the relative position of cortex (Ctx), striatum (Str), and thalamus (Th) in adult mouse brain.

(D) Violin plots indicating the module scores of glial clusters (x-axis) for each glial cell type (one per plot).

(E) UMAP plot of dorsal striatum data with color indicating expression of the medium spiny neuron (MSN) markers *Rgs9* and *Ppp1r1b*. Clusters with high levels of both markers were further analyzed in (F).

(F) Dot plot showing normalized expression of transcripts previously shown to be enriched in D1-, D2-, matrix, patch, and eccentric (Ecc) MSNs (y-axis) in each MSN cluster identified in the present study (x-axis).

(G) UMAP plot with color indicating normalized *Pvalb* RNA level.

(H) UMAP plot with color indicating normalized *Cck* RNA level.

(I) UMAP plot with color indicating normalized *Pthlh* RNA level.

(J) UMAP plot with color indicating normalized *Sst* and *Npy* RNA levels.

(K) Heatmap showing normalized average expression of genes within the *Df(16)1/+* hemizygous region. X-axis indicates mouse ID and genotype. All nuclei except those shown to be contaminants from the cortex and thalamus in (B) and (C) were combined to generate the heatmap.

(L) UMAP plot with color indicating genotype at the *Df(16)1* locus.

(M) Bar plot showing the number of nuclei within each cluster. Color indicates genotype at the *Df(16)1* locus. Data in (B) were produced by snRNA-seq analysis of 43,319 nuclei representing all nuclei that passed quality control parameters, including some from the thalamus and cortex.

Data in (C–M) were produced by snRNA-seq analysis of 35,167 nuclei derived from the dorsal striatum. expression: normalized average gene expression; proportion: proportion of cells expressing a marker within a cluster.

**Figure S5. Additional behavioral measures in 22q11DS mice (Related to Figure 7).**

(A) There was no difference in the number of lever presses (left) or rewards earned (right) in the RR task between vehicle- and C21-injected *Df(16)1/+* male mice (paired *t* test for lever presses, *p* = 0.12; for rewards earned, *p* = 0.23).

(B) C21 injection decreased performance on the FR1 task in *Df(16)1/+* male mice (paired *t* test, \**p* = 0.03).

(C) *Df(16)1/+* mice preferred to consume sweetened condensed milk (SCM) over food chow when both were freely available after injections of vehicle and C21 (three-way ANOVA, \*\*\*\**p* < 0.0001). There was no difference in the amount of SCM consumed based on injection type (Tukey’s multiple comparison, *p* = 0.98).

(D) There was no difference in the number of lever presses (left) or rewards earned (right) in the RR task between vehicle- and C21-injected *Df(16)1/+* mice (two-way RM ANOVA for lever presses, *p* = 0.14; for rewards earned, *p* = 0.24). There was a sex difference in RR performance (two-way RM ANOVA, \**p* = 0.02).

(E) There was no difference in performance on the FR1 task after C21 injection (two-way RM ANOVA, *p* = 0.16).

(F) There was no difference in the amount of freely available rodent chow consumed in the FR1 choice task after injection of C21 (two-way RM ANOVA, *p* = 0.08).

(G) Mice preferred SCM over standard rodent chow when both were freely available regardless of injection type (three-way ANOVA, \*\*\*\**p* < 0.0001). There was no difference in SCM consumption based on injection type (Tukey’s multiple comparison, *p* = 0.73).

(H) There was no difference in the number of lever presses (left) or rewards earned (right) in the RR task between vehicle- and C21-injected *Df(16)1/+* mice (two-way RM ANOVA for lever presses, *p* = 0.28; for rewards earned, *p* = 0.3).

(I) There was no difference in performance on the FR1 task after C21 injection (two-way RM ANOVA, *p* = 0.1).

(J) There was no difference in the consumption of freely available rodent chow in the FR1 choice task after injection of C21 (two-way RM ANOVA, *p* = 0.57).

(K) Mice preferred SCM over standard rodent chow when both were freely available regardless of injection type (three-way ANOVA, \*\*\*\**p* < 0.0001). There was no difference in SCM consumption based on injection type (Tukey’s multiple comparison test, *p* = 0.99).

Data shown are mean ± SEM with individual data points overlaid in (A–K).

Unless noted, there were no sex differences.

## STAR Methods

### Key Resources Table

| REAGENT or RESOURCE | SOURCE | IDENTIFIER |
| --- | --- | --- |
| Bacterial and virus strains |  |  |
| pAAV5-hSyn-hChR2(H134R)-EYFP | Addgene | #26973-AAV5 |
| pENN-AAVrg-hSyn-HI-eGFP-Cre.wpre.SV40 | Addgene | #105540-AAVrg |
| pAAV-hSyn-DIO-hM4D(Gi)-mCitrine | Addgene | #50455-AAV8 |
| pAAV-hSyn-DIO-hM4D(Gi)-mCherry | Addgene | #44362-AAV5 |
| pAAV-hSyn-DIO-HA-hM3D(Gq)-IRES-mCitrine | Addgene | #50454-AAV8 |
| pAAV-hSyn-GRAB-ACh3.0 | Addgene | #121922-AAV9 |
| Chemicals, peptides, and recombinant proteins |  |  |
| SR 95531 hydrobromide | Tocris Bioscience | Cat #1262 |
| CGP 55845 hydrochloride | Tocris Bioscience | Cat #1248 |
| AP5 | Tocris Bioscience | Cat #0106 |
| DNQX | Tocris Bioscience | Cat #018910 |
| TTX | Tocris Bioscience | Cat #1069 |
| Actinomycin D | Sigma | Cat #A1410 |
| DREADD agonist 21 dihydrochloride<br>(Compound 21) | Tocris Bioscience | Cat #6422 |
| AF-DX-116 | Tocris Bioscience | Cat #1105 |
| Deposited data |  |  |
| snRNA-seq | This paper | GEO: GSE293229 |
| R code for snRNA-seq analysis | This paper | Github:<br>https://github.com/ZakharenkoLab/Df1_dorsalstriatum_snRNAseq |
| Experimental models: Organisms/strains |  |  |
| *Df(16)1/+* | Lindsay et al., (1999)<sup>74</sup> | N/A |
| ChAT<sup>Cre</sup> | Jax Labs | Strain #006410 |
| Ai14 | Jax Labs | Strain #007914 |
| Software and algorithms |  |  |
| pClamp | Molecular Devices | RRID:SCR_011323 |
| PrairieView | Bruker | N/A |
| Prism | GraphPad | RRID:SCR_002798 |
| MiniAnalysis | Synaptosoft | N/A |
| Fiji | Schindelin et al., (2012)<sup>123</sup> | RRID:SCR_002285 |
| MED PC | Med Associates | RRID:SCR_012156 |
| TopScan | CleverSys | RRID:SCR_017141 |
| BZ-X Viewer | Keyence | RRID:SCR_016979 |
| RStudio v4.3.1 | Posit | RRID:SCR_000432 |
| Star v2.7 | Dobin et al.<sup>124</sup> | RRID: SCR_004463 |
| RSEM v1.3.1 | Li et al.<sup>125</sup> | RRID: SCR_000262 |
| VoxHunt v1.1.0 | Fleck et al.<sup>126</sup> | RRID: SCR_023829 |
| g:Profiler | Raudvere et al.<sup>127</sup> | RRID: SCR_006809 |
| ggplot2 | Wickham<sup>128</sup> | RRID: SCR_014601 |
| ComplexHeatmap v2.10.0 | Gu; Gu et al.<sup>129,130</sup> | RRID: SCR_017270 |
| bcl2fastq v2.20.0.422 | Illumina | RRID: SCR_015058 |
| CellRanger v7.0.0 | 10xGenomics | RRID: SCR_017344 |
| SoupX v1.6.2 | Young and Behjati<sup>131</sup> | RRID: SCR_019193 |
| Seurat v4.4.0 | Satija Lab<sup>132–135</sup> | RRID: SCR_016341 |
| Harmony v 1.2.0 | Korsunsky et al.<sup>136</sup> | RRID: SCR_022206 |
| DESeq2 v1.42.1 | Love et al.<sup>137</sup> | RRID: SCR_015687 |
| clusterProfiler v4.12.6 | Yu et al.<sup>138</sup> | RRID: SCR_016884 |

### Animals

All experiments were carried out in accordance with the United States Public Health Service *Guide for Care and Use of Laboratory Animals*. Experiments were approved by the Institutional Animal Care and Use and Committee at St. Jude Children’s Research Hospital.

Female and male mice between 2 and 8 months old were used for experiments. The generation of *Df(16)1/+*, ChAT^Cre^, and Ai14 mice was as previously reported.^74,139,140^ *Df(16)1/+* mice were backcrossed onto the C57Bl/6J genetic background for at least 10 generations. Littermate mice were housed 2-5 per cage under a regular light/dark cycle (lights on at 0600h, off at 1800h) with *ad libitum* access to food and water, except during operant training.

### Method Details

#### Surgical Procedures

Adult mice (>2 months old) were anesthetized with 2.5% isoflurane in 100% O_2_ and mounted in a stereotaxic setup. Anesthesia was maintained with 1.5% isoflurane for the duration of the surgery. Under aseptic conditions, a midline incision was made, and burr holes were drilled bilaterally above the Pf (coordinates in mm from bregma: AP –2.2, ML ±0.5, DV –3.5 from top of brain) or the DMS (coordinates in mm from bregma: AP +0.5, ML ±2.0, DV –2.4 from top of brain). A 32-gauge needle was inserted to pressure inject 300 or 500 nl of virus (to the Pf and DMS, respectively) with a pump rate of 25 nl/min. After surgery, mice were returned to their home cages for 3 or 8 weeks to allow for virus expression within the DMS or the Pf-DMS pathway, respectively.

The following viruses were purchased from Addgene and used for intracranial infusions: pAAV5-hSyn-hChR2(H134R)-EYFP (Addgene #26973-AAV5), pENN-AAVrg-hSyn-HI-eGFP-Cre.wpre.SV40 (Addgene #105540-AAVrg), pAAV-hSyn-DIO-hM4D(Gi)-mCherry (Addgene #44362-AAV5), pAAV-hSyn-DIO-hM4D(Gi)-mCitrine (Addgene #50455-AAV8), pAAV-hSyn-DIO-HA-hM3D(Gq)-IRES-mCitrine (Addgene #50454-AAV8), and pAAV-hSyn-GRAB-ACh3.0 (Addgene #121922-AAV9).

#### Behavioral Tests

##### Food Restriction

Mice undergoing operant training were food restricted at 70%-85% ad libitum of their daily intake to maintain ∼20% body mass reduction.^141^ Food intake during training was measured, and food was made available for 2 h after each session to ensure that each mouse was adequately fed.

##### Feeding Tests

To decrease the likelihood of neophobia obscuring a potential food-type preference, we conducted two 1-h–long exposures of the mice to SCM in standard housing cages on consecutive days. Mice were then tested for food-type preference. For 1 h, mice were given the choice between standard pelleted rodent chow and SCM. Both food types were weighed before and after the trials to calculate the amount ingested.

For SCM consumption experiments, mice were given 1 h access to SCM in standard housing cages. SCM was weighed before and after the trial to calculate the amount ingested.

##### Operant Behavior

The operant behavior test was performed as previously described.^142,143^

##### Part 1. Trough acclimation, presentation, and lever pressing training

All operant tasks were performed in operant conditioning chambers (MED-307A, Med Associates Inc.), with training and reward schedule trials each lasting 60 min unless otherwise stated. On the first day, mice were placed into an operant conditioning chamber and trained to obtain SCM from the feeding trough in two stages. During the initial stage, mice received SCM each time they entered the food trough. This stage was completed after 20 reinforced entries. The second stage had a variable time schedule in which a dipper cup containing SCM was raised into the feeding trough and the mouse was given 10 s to obtain the reward. Mice remained at the trough training stage until at least 20 entries of 30 presentations were made on 2 consecutive days. All reinforcer deliveries were accompanied by an 8 kHz tone.

For lever pressing training, mice were placed into the operant conditioning chamber with access to a lever that, when pressed, gave access to SCM for 8 s. To complete this stage, mice were required to obtain 50 rewards on 2 consecutive days. The total number of days it took mice to complete trough and lever training was calculated and presented as Days to Criterion. If progress was not made on Part 1 after 10 days, the mouse was eliminated from the study.

##### Part 2. Fixed Ratio 1 (FR1) Choice Task

To test the willingness of mice to work for a more rewarding food type (i.e., SCM) in the presence of a freely available, less-preferred food type (i.e., chow), we placed the mice into the operant chamber with *ad libitum* access to chow and water, and the criterion to earn SCM was 1 lever press. The number of lever presses and the amount of standard chow consumed were measured.

##### Part 3. Random Ratio (RR) Task

On a separate day, mice were placed in the operant chamber with the criterion to earn SCM set to vary randomly during the session, with the average number of lever presses over the course of the experiment set to 10. The number of lever presses and rewards earned were measured.

##### Part 4. Progressive Ratio +1 (PR+1) and Progressive Ratio +4 (PR+4) Tasks

To assess the motivation to earn a rewarding food, we used a PR model.^19,78,79^ Two ratios that vary in work required were used: PR+1 and PR+4. In PR+1, the initial lever press criterion was set at 1 and increased by 1 after each trial (i.e., 1, 2, 3, 4). In PR+4, the initial lever press criterion was set at 4 and increased by 4 after each trial (i.e., 4, 8, 12, 16). Trials lasted for 60 min (PR+1) or 90 min (PR+4) or were terminated if the mouse did not lever press for 5 min. The number of lever presses and rewards earned were recorded.

##### C21 Injection

For *in vivo* DREADDs experiments, the above training schedule was followed, with Parts 2– 4 and food preference tests repeated on separate days to account for C21 and vehicle (saline) injections. Saline and C21 (10 mg/kg at 1 mg/ml) were injected intraperitoneally 30 min before the start of the session. The order of injection type was randomized and counterbalanced. The experimenter was blinded to the experimental condition.

##### Open-Field Test

Mice were acclimated to the testing room for 1 h prior to testing. The open field arena consisted of four clear plastic walls and a blue matte floor, with a partition between the arenas, creating a 16 × 16-in cube with 14-in walls and no top. Mice were recorded from above using a Panasonic camera (WV-CP304) run into a Windows-based computer via an Everplex 4CQ (EverFocus Electronics Corporation) and controlled remotely by WinTV-HVR-1955 (Hauppauge!). Video capture and analysis was performed using TopScan Real Time Option (v3.0, CleverSys Inc.). Each mouse was recorded for 30 min then returned to their home cage. The arena was cleaned after each trial with 70% ethanol and a disposable cloth and allowed to dry for 10 min.

#### Acute Slice Preparation

After rapid decapitation, mouse brains were removed and submerged in ice cold cutting solution (in mM: 125 choline-Cl, 2.5 KCl, 0.4 CaCl_2_, 6 MgCl_2_, 1.25 NaH_2_PO_4_, 26 NaHCO_3_, and 20 D-glucose, 300–310 mOsm, under 95% O_2_/5% CO_2_). Coronal or oblique horizontal slices of 250-µm thickness were made of the DMS, nucleus accumbens shell, or Pf. Oblique horizontal slices of 250-µm thickness were made to preserve Pf projections to the DMS.^55^ Slices were incubated in ACSF (in mM: 125 NaCl, 2.5 KCl, 2 CaCl_2_, 2 MgCl_2_, 1.25 NaH_2_PO_4_, 26 NaHCO_3_, 20 D-glucose, 300–310 mOsm, with 95% O_2_/5% CO_2_) for 30 min before being stored at room temperature for the remainder of the day.

#### Whole-Cell Patch-Clamp Electrophysiology

Tissue slices were placed in a recording chamber mounted on a two-photon laser scanning microscope (Bruker), and superfused with ACSF at 32°C ± 1°C (2–3 ml/min). Cells were visualized under two-photon guidance using PrairieView v5.5 software. Whole-cell recordings were made from visually identified MSNs, CHIs, and Pf neurons by using borosilicate glass pipettes (Sutter, open pipette resistance of 3–6 MΩ). For current-clamp recordings, glass pipettes were filled with an internal pipette solution containing the following (in mM): 115 potassium gluconate, 20 KCl, 10 HEPES, 4 MgCl_2_, 0.1 EGTA, 4 ATP-Mg_2_, 0.4 GTP-Na, and 10 creatine phosphate-Na_2_, pH 7.4, 290–295 mOsm. For voltage-clamp recordings, pipettes were filled with the following (in mM): 125 CsMeSO_3_, 2 CsCl, 10 HEPES, 0.1 EGTA, 4 ATP-Mg_2_, 0.3 GTP-Na, 10 creatine phosphate-Na_2_, 5 QX-314, and 5 TEA-Cl, pH 7.4, 290-295 mOsm. Neurons were voltage clamped at –60 mV with a MultiClamp 700B amplifier (Molecular Devices). Signals were digitized at 20 kHz with an Axon Digidata 1550B (Axon Instruments) and filtered at 2 kHz with Clampex 10.7 software. The liquid junction potential was calculated to be –10 mV and was corrected for in each recording.

Synaptic transmission was assessed using paired electrical or optical stimulation (2-ms pulse duration at 10 Hz) delivered every 20 s for the duration of the experiment. Electrically evoked EPSCs were elicited using a bipolar concentric stimulating electrode (World Precision Instruments) placed in the Pf of oblique horizontal slices or locally within the shell of the nucleus accumbens connected to a stimulus-isolation unit (Iso-Flex, AMPI). Optically evoked EPSCs were elicited by activating Pf terminals with LED blue light pulses via a fiber optic cable (473 nm, M470F3, Thor Labs) placed locally within the DMS approximately 100 µm from the recorded neuron. EPSCs (evoked and spontaneous) were recorded in the presence of the GABA_A_ and GABA_B_ receptor antagonists, SR 95531 (1 µM) and CGP 55845 (1 µM). To ensure accurate stimulation delivery across input-output experiments, we measured the impedance of the stimulating electrode and the optical output of the fiber optic regularly, and the electrode or optical fiber were replaced if the measurement deviated by more than 10%. Access resistance of the recording electrode was measured throughout the recording, and cells were discarded from analysis if the resistance changed by more than 15%.

To measure input resistance and evoked firing rates, we injected a series of hyperpolarizing and depolarizing step currents into cells in current-clamp mode (+10-pA steps from –50 pA up to +240 pA). The rheobase was measured by the injection of a depolarizing ramp current (from –20 pA to +200 pA) into cells in current-clamp mode. The current at which the first AP was generated was recorded. To determine tonic activity in CHIs, we recorded the activity state of the cell immediately (within 1 min) upon entering whole-cell mode.

#### GRAB-ACh Imaging

Three weeks after viral injection of pAAV-hSyn-GRAB-ACh3.0^97^ into the DMS, coronal slices were made and placed in a recording chamber under a two-photon laser-scanning microscope. Two-photon laser-scanning microscopy of GRAB-ACh was performed using an Ultima imaging system (Bruker), a Ti:sapphire Chameleon Ultra femtosecond-pulsed laser (Coherent, 930 nm), and a 60× (NA 0.9) water-immersion infrared objective (Olympus). Regions with high GRAB-ACh expression were imaged for 2.5 min with a Galvo scanner at a rate of 1 frame/s with a field of view of 512 × 512 µm. Electrical stimulation was delivered locally (approximately 250 µm away from the imaging location) at 10, 20, 50, and 100 Hz (10 pulses at the desired frequency delivered 10 times every 1.5 s). Electrical stimulation was set at 300 µA for all imaging experiments. GRAB-ACh signals were processed from the video sequences by extracting the fluorescence intensities of the entire imaging field over time. Activity signals were normalized to baseline levels, and peak amplitudes were calculated as dF/F, where F is the baseline fluorescence intensity. Calculated dF/F values were imported into the data analysis software, Clampfit 10.7, to detect the average peak amplitude after stimulation. Following the stimulation protocol, a new slice was selected. Stimulation frequency delivery was randomized across slices.

#### Immunofluorescence and Histologic Verification

Mice expressing the tdTomato fluorescent protein under the ChAT promoter were deeply anesthetized by intraperitoneal injection of Avertin (0.2 ml/10 g, 1.25% v/v solution) and transcardially perfused with phosphate buffered solution (PBS) followed by 4% paraformaldehyde (PFA) in 0.1 M phosphate buffer. The brains were fixed in PFA overnight, then washed in PBS for 25 min with shaking, protected from light. The brains were blocked and mounted in 4% agarose. 50-µm thick coronal serial slices were obtained using a Leica VT1000. After being mounted and coverslipped, slides were imaged using a Keyence BZ-X700 or a BZ-X800 with a 4× objective lens (0.13 NA, 0.2 NA, respectively). Images were stitched in BZ-X Viewer software to produce a whole coronal brain section.

For automatic detection of tdTomato^+^ CHIs, the StarDist^144^ method for nucleus segmentation was used as a Fiji plug-in.^123^ After selecting the dorsal striatum in each image in Fiji, we used the pretrained StarDist model ‘Versatile (fluorescent nuclei)’ with the default method parameters to detect tdTomato^+^ cells.

#### Single-Nucleus RNA Sequencing

##### Sample Collection

Transcardial perfusion was performed on four 2-month-old *Df(16)1/+* and four WT littermate mice of both sexes with ice cold, oxygenated ACSF consisting of the following (in mM): 125 NaCl, 2.5 KCl, 1.25 NaH_2_PO_4_, 26 NaHCO_3_, 25 glucose, 5 ethyl pyruvate, 0.4 sodium ascorbate, 1 MgCl_2_, and 2 CaCl_2_. The ACSF also contained the following blockers of neuronal activity and transcription (in µM): 20 AP5, 20 DNQX, 0.1 TTX, 5 Actinomycin D. The samples were quickly extracted, flash frozen in liquid nitrogen, and stored at –80°C until nuclear dissociation.

##### Nuclei Dissociation

Nuclei dissociation was performed as described.^145,146^ Briefly, frozen tissue was mechanically dissociated with a Dounce homogenizer in detergent lysis buffer containing 0.32 M sucrose, 10 mM HEPES (pH 8.0), 5 mM CaCl_2_, 3 mM magnesium acetate, 0.1 mM EDTA, 1 mM DTT, and 0.1% Triton-X100. The resulting homogenate was filtered through a 40 µm strainer and washed with the same solution described without the Triton-X100 added. Nuclei were then centrifuged at 3200×*g* for 10 min at 4°C and the supernatant was decanted. A sucrose dense solution containing 1 M sucrose, 10 mM HEPES (pH 8.0), 3 mM magnesium acetate, and 1 mM DTT was carefully layered underneath the remaining supernatant and then spun at 3200×*g* for 20 min at 4°C. The supernatant was discarded, and the final remaining nuclei resuspended in 0.4 mg/ml BSA and 0.2 U/µl Lucigen RNAse inhibitor diluted in PBS. Between 5,000 and 10,000 nuclei were inspected and counted on a hemacytometer before loading on to the 10X chromium controller (10X Genomics Catalog #1000171).

##### Library Preparation and Sequencing

Nuclei were loaded onto the 10X Genomics Chromium Controller using the Next GEM Single Cell 3’ Reagents Kit Version 3.1 (10X Genomics Cat #1000121) Dual Index (10X Genomics Cat #PN-1000215) gene expression profiling assay according to the manufacturer’s instructions. After capture and lysis, cDNA synthesis, amplification, and library construction were performed on an ABI ProFlex PCR System (Applied Biosystems). The libraries were quantified with the Quant-iT Pico Green dsDNA assay (ThermoFisher) and through low-pass sequencing with a MiSeq nano kit (Illumina). Quality assessments were conducted on the Agilent Tapestation (Agilent Cat #G2964AA) with D5000 and D1000 reagents. Paired-end 100-cycle sequencing (Read length configuration: 28-10-10-90) was carried out on a NovaSeq 6000 (Illumina) in the St. Jude Hartwell Center Genome Sequencing Core.

#### Statistical Analysis

All data are presented as mean ± SEM with individual data points overlaid on bar graphs. All data were analyzed with GraphPad Prism v10.2.2, unless otherwise noted.

##### Behavioral Data Analysis

All behavioral data were analyzed offline with GraphPad Prism v10.2.2. Lever presses and rewards earned on all operant tasks were compared using the following statistical measures.

Comparisons between genotypes were made using sex (female/male) x genotype (WT/*Df(16)1/+*) ANOVAs. Comparisons for DREADDs experiments were made using sex x drug (vehicle/C21) repeated measures (RM) ANOVAs. To quantify the difference in PR performance between experimental groups, we calculated difference ratios. The average number of lever presses made by *Df(16)1/+* mice (Y) were subtracted and divided by the average number of lever presses made by WT mice (X): (Y-X)/X. To calculate the difference ratios between C21 and vehicle injections, the same formula was used on individual mice with Y equaling lever presses made following C21 injection and X equaling lever presses with vehicle injection. Difference ratio scores were compared against a theoretical mean of 0 using a one-sample *t* test.

Comparisons of overall SCM consumption, the number of days it took mice to reach training criterion, the overall time spent in the center of the open field, and the overall distance traveled in the open field were made using sex x genotype ANOVAs. Comparisons on food preference tests were made using sex x drug x food type (SCM/chow) ANOVAs or mixed-design ANOVAs (sex x genotype x food type). To compare cumulative distance traveled and the time spent in the center of the open field arena across time, time x genotype mixed-design ANOVAs were used.

##### Electrophysiology Data Analysis

All electrophysiologic experiments were analyzed offline using Clampfit 10.7 software and statistical comparisons were conducted in GraphPad Prism v10.2.2. All raw EPSCs amplitudes were measured and averaged per minute. For input-output experiments, averaged EPSC amplitudes were plotted. Differences between groups across stimulus intensities were assessed using a mixed-design ANOVA (stimulus intensity x genotype). A mixed-design ANOVA (current intensity x genotype) was used to compare evoked firing rates across multiple step current injections between genotypes. Differences in membrane properties between genotypes were analyzed using unpaired *t* tests.

To measure changes in synaptic transmission as a response to drug application, we expressed EPSC amplitudes as a percentage change from baseline (min 1–5). To determine a difference in synaptic transmission following drug application, we compared the average of the last 5 min across groups by using an unpaired *t* test. PPR was calculated by dividing the peak amplitude of EPSC2 by the peak amplitude of EPSC1. Differences in PPR were compared between groups using an unpaired *t* test.

The amplitude and frequency (presented as interevent interval) of sEPSC were detected and calculated by MiniAnalysis v6.0.7 (SynaptoSoft) and compared using unpaired *t* tests. The distribution of tonically active versus silent CHIs was compared using a Chi-square test.

##### GRAB-ACh Imaging Analysis

Peak GRAB-ACh dF/F amplitudes were averaged across slices, and differences between genotypes were compared using a mixed-design ANOVA (stimulation frequency x genotype).

##### snRNA-seq Data Processing and Analyses

Sequencing reads were processed using 10X Genomics Cell Ranger v7.0.0, with mouse reference genome mm10-2020-A provided by 10X Genomics. Cell Ranger was run with –force-cells 8,000 flag. SoupX v1.6.2 was run to generate adjusted count matrices with corrections for ambient RNA contamination applied. Quality control filtering, clustering, dimensionality reduction, visualization, and differential gene expression were performed using Seurat v4.4.0 with R v4.3.1. Each dataset was filtered so that genes expressed in at least three or more cells were included in the final dataset. Cells were excluded if they (1) expressed greater than 1% of mitochondrial genes, (2) had fewer than 1,000 genes expressed, or (3) had a unique molecular identifier count below 500. Datasets were individually log-normalized using Seurat’s NormalizeData with default parameters. Cell cycle scoring was conducted using the associated S and G2M phase gene list from Tirosh, et al.^147^ with the CellCycleScoring command in Seurat. We calculated 3,000 features that exhibit high cell-to-cell variation in the dataset using Seurat’s FindVariableFeature function. Next, we scaled the data by linear regression against the number of reads using Seurat’s ScaleData function with default parameters. The variable genes were projected onto a low-dimensional subspace by using principal component analysis through Seurat’s RunPCA function with default parameters. The number of principal components (Npcs) was selected based on inspection of the plot of variance explained (Npcs = 30).

Datasets were integrated using Harmony (v1.2.0) integration with default parameters.^136^ A shared-nearest-neighbor graph was constructed based on the Euclidean distance in the low-dimensional subspace by using Seurat’s FindNeighbors with dims = 1:30 and default parameters. Integrated datasets then underwent non-linear dimensional reduction and visualization through Uniform Manifold Approximation and Projection. Clusters were identified using a resolution of 0.2 and the Leiden algorithm for the Harmony integrated datasets. Differentially expressed genes for each cluster were identified using Seurat’s FindAllMarkers function (parameters: logfc.threshold = 0.25, test.use = “wilcox”, min.pct = 0.1, return.thresh = 0.01). Differentially expressed genes for comparing *Df(16)1/+* and WTs within each cluster were generated using DESeq2 and pseudobulk expression. The pseudobulk expression matrix was generated using Seurat’s AggregateExpression function to group expression based on cluster, sample ID, and condition.

Individual clusters defined with data processed using Harmony (see above) were evaluated for known cell type markers based on the differentially expressed genes identified for each cluster (see above). Additionally, module scores for glia/non-neuronal cell types (i.e. endothelial cells, astrocytes, microglia, oligodendrocytes, and oligodendrocyte precursor cells) were calculated for each cell by using Seurat’s AddModuleScore function with lists of cell type markers (**Table S1**) for each cell type (parameters features = list(gene.markers), name = cell.type). Brain structure marker heatmaps were generated using VoxHunt v1.1.0 with regional markers from the *Allen Developing Mouse Brain Atlas* P56 data set. GO overrepresentation analysis of differentially expressed genes (*p* < 0.01) in CHIs was performed using the enrichGO function from clusterProfiler v4.12.6 with default parameters for mouse genes (OrgDb = org.Mm.eg.db) and cellular component terms (ont = “CC”).

## Supplemental Tables

**Table S1.** Genes used to calculate module scores for glial cell types.

| Cell type | Gene |
| --- | --- |
| Endothelial Cell | Vwf |
| Endothelial Cell | Cldn5 |
| Endothelial Cell | Abcb1 |
| Endothelial Cell | Ebf1 |
| Endothelial Cell | Itm2a |
| Endothelial Cell | Tie1 |
| Endothelial Cell | Flt1 |
| Endothelial Cell | Notch3 |
| Endothelial Cell | Cxcl12 |
| Endothelial Cell | Pecam1 |
| Endothelial Cell | Foxq1 |
| Endothelial Cell | Tbx3 |
| Endothelial Cell | Cavin2 |
| Endothelial Cell | Cp |
| Endothelial Cell | Mgp |
| Endothelial Cell | Tm4sf1 |
| Endothelial Cell | Slc38a5 |
| Endothelial Cell | Anpep |
| Endothelial Cell | Itgb1 |
| Endothelial Cell | Cd34 |
| Endothelial Cell | Cd93 |
| Endothelial Cell | Icam2 |
| Endothelial Cell | Eng |
| Endothelial Cell | Il1r1 |
| Endothelial Cell | Prom1 |
| Endothelial Cell | Thbd |
| Endothelial Cell | Cdh5 |
| Endothelial Cell | Bsg |
| Endothelial Cell | Cd151 |
| Endothelial Cell | Cd248 |
| Endothelial Cell | Kdr |
| Endothelial Cell | Emcn |
| Endothelial Cell | Esam |
| Endothelial Cell | Itga4 |
| Endothelial Cell | Klf4 |
| Endothelial Cell | Podxl |
| Endothelial Cell | Thsd1 |
| Endothelial Cell | Tie1 |
| Endothelial Cell | Egfl7 |
| Astrocyte | Aldh1l1 |
| Astrocyte | Gja1 |
| Astrocyte | Adgrv1 |
| Astrocyte | Gpc5 |
| Astrocyte | Aqp4 |
| Astrocyte | Sdc4 |
| Astrocyte | Aldh1a1 |
| Astrocyte | Sox9 |
| Astrocyte | Slc1a2 |
| Astrocyte | Tnc |
| Astrocyte | Gfap |
| Astrocyte | Slc1a3 |
| Astrocyte | Slc1a2 |
| Astrocyte | Glul |
| Astrocyte | Slc38a3 |
| Astrocyte | Gldc |
| Astrocyte | Hibadh |
| Astrocyte | Serhl |
| Astrocyte | Prodh |
| Astrocyte | Apoe |
| Astrocyte | Eci1 |
| Astrocyte | Eci2 |
| Astrocyte | Gk |
| Astrocyte | Cpt1a |
| Microglia | Cx3cr1 |
| Microglia | Csf1r |
| Microglia | Tmem119 |
| Microglia | Itgam |
| Microglia | C1qb |
| Microglia | Ly86 |
| Microglia | Runx1 |
| Microglia | P2ry12 |
| Microglia | Fcrls |
| Microglia | Entpd1 |
| Microglia | Siglech |
| Microglia | Ccr5 |
| Microglia | Cd33 |
| Microglia | Gpr34 |
| Microglia | Tgfb1 |
| Microglia | Tgfbr1 |
| Microglia | Serinc3 |
| Microglia | Mertk |
| Microglia | Spi1 |
| Microglia | Mef2a |
| Microglia | Ctsd |
| Microglia | Trem2 |
| Oligodendrocyte | Mbp |
| Oligodendrocyte | Mog |
| Oligodendrocyte | Mobp |
| Oligodendrocyte | Plp1 |
| Oligodendrocyte | Olig2 |
| Oligodendrocyte | Cldn11 |
| Oligodendrocyte | Tspan2 |
| Oligodendrocyte | Serinc5 |
| Oligodendrocyte | Opalin |
| Oligodendrocyte | Mal |
| Oligodendrocyte | Dock5 |
| Oligodendrocyte | Hapln2 |
| Oligodendrocyte | Apod |
| Oligodendrocyte | Aspa |
| Oligodendrocyte | Carns1 |
| Oligodendrocyte | Nkx6-2 |
| Oligodendrocyte | Cercam |
| Oligodendrocyte | Tfeb |
| Oligodendrocyte | Klk6 |
| Oligodendrocyte | Pmp22 |
| Oligodendrocyte Precursor Cell | Cspg4 |
| Oligodendrocyte Precursor Cell | Pdgfra |
| Oligodendrocyte Precursor Cell | Tcf7l2 |
| Oligodendrocyte Precursor Cell | Ptprz1 |
| Oligodendrocyte Precursor Cell | C1ql1 |
| Oligodendrocyte Precursor Cell | Vcan |
| Oligodendrocyte Precursor Cell | Bmp4 |
| Oligodendrocyte Precursor Cell | Neu4 |
| Oligodendrocyte Precursor Cell | Tnr |
| Oligodendrocyte Precursor Cell | Tns3 |
| Oligodendrocyte Precursor Cell | Fyn |
| Oligodendrocyte Precursor Cell | Itpr2 |
| Oligodendrocyte Precursor Cell | Bcas1 |
| Oligodendrocyte Precursor Cell | Rras |
| Oligodendrocyte Precursor Cell | Cnksr3 |
| Oligodendrocyte Precursor Cell | Gpr17 |
| Oligodendrocyte Precursor Cell | Tmem100 |

**Table S2.**
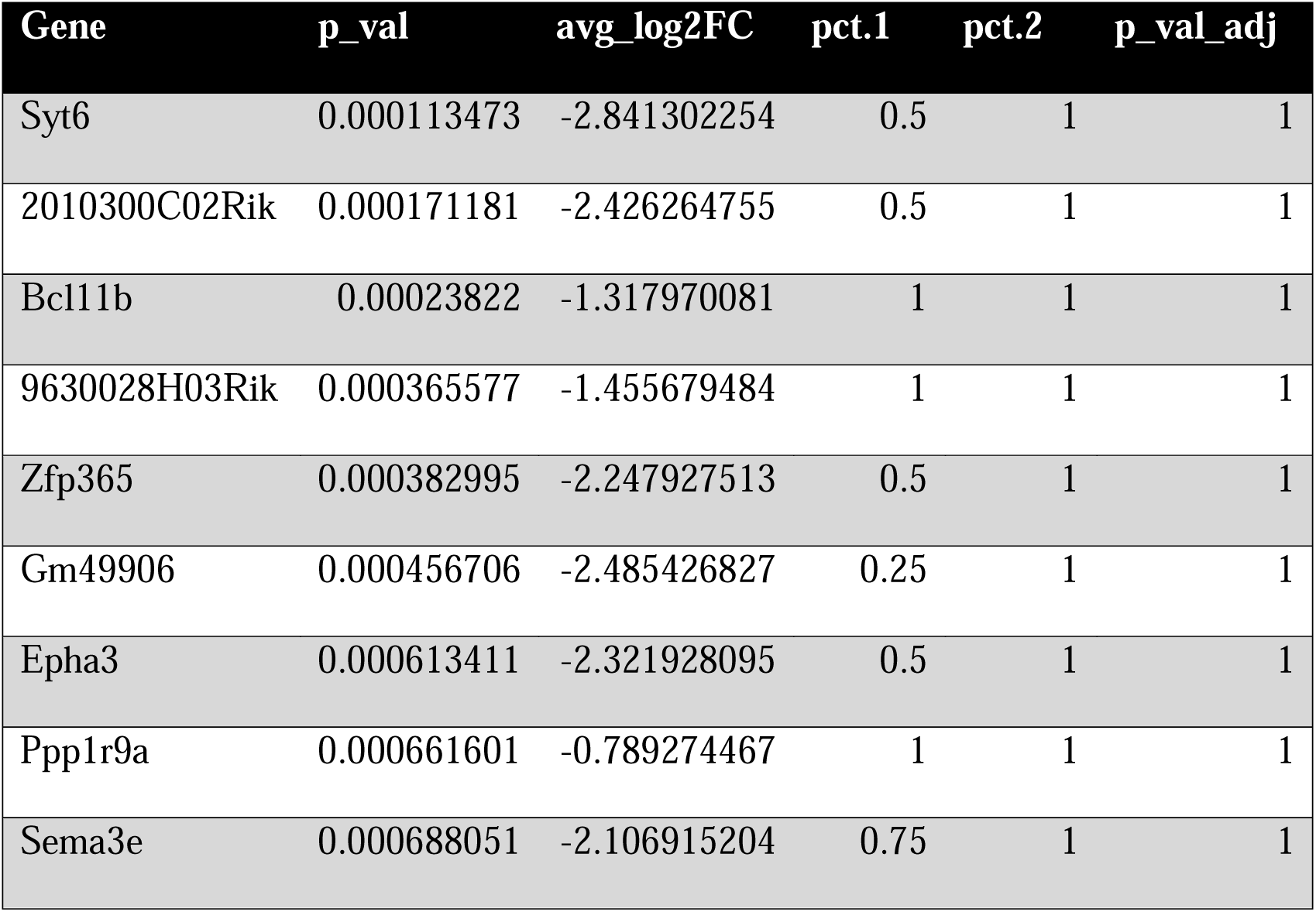
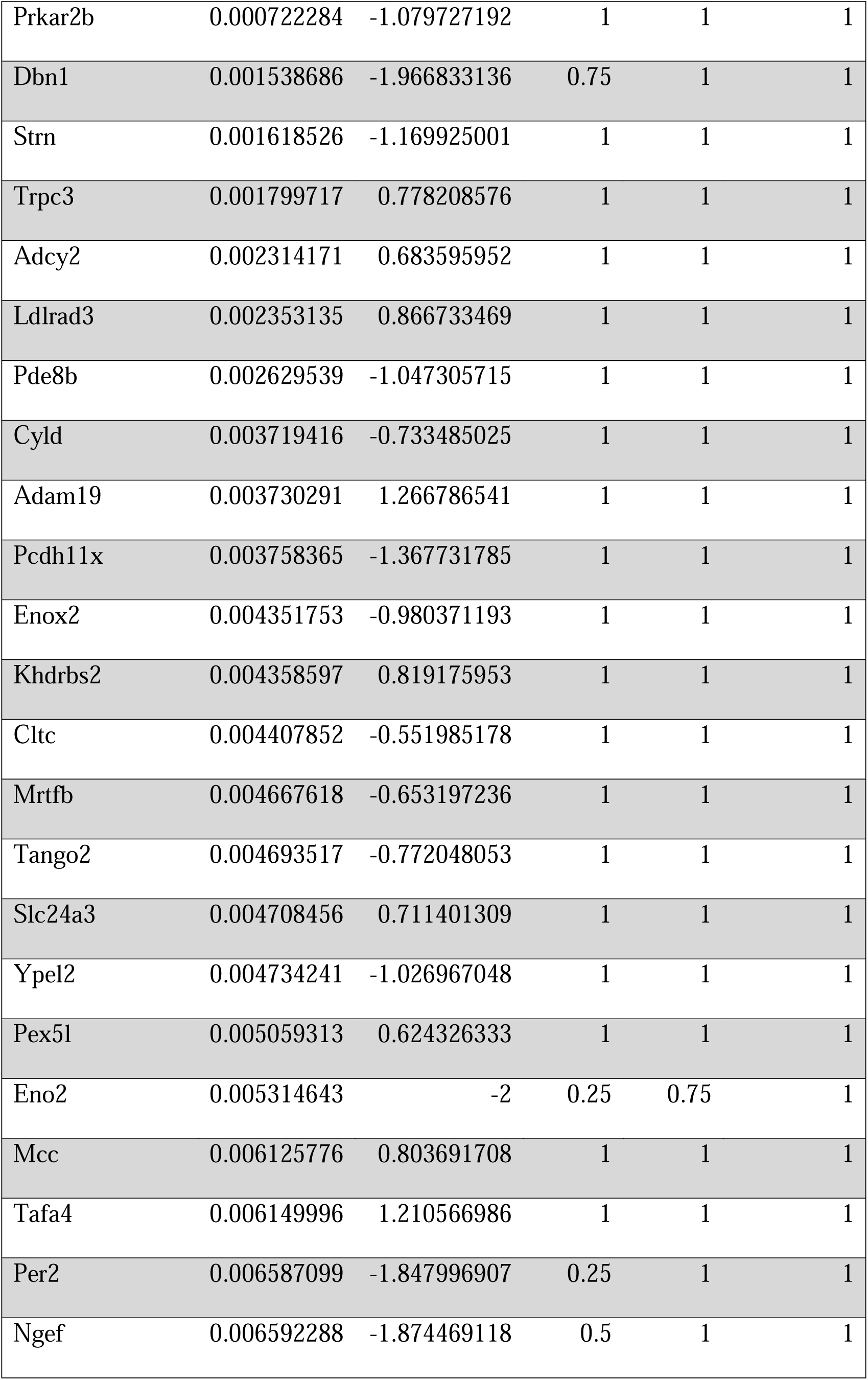
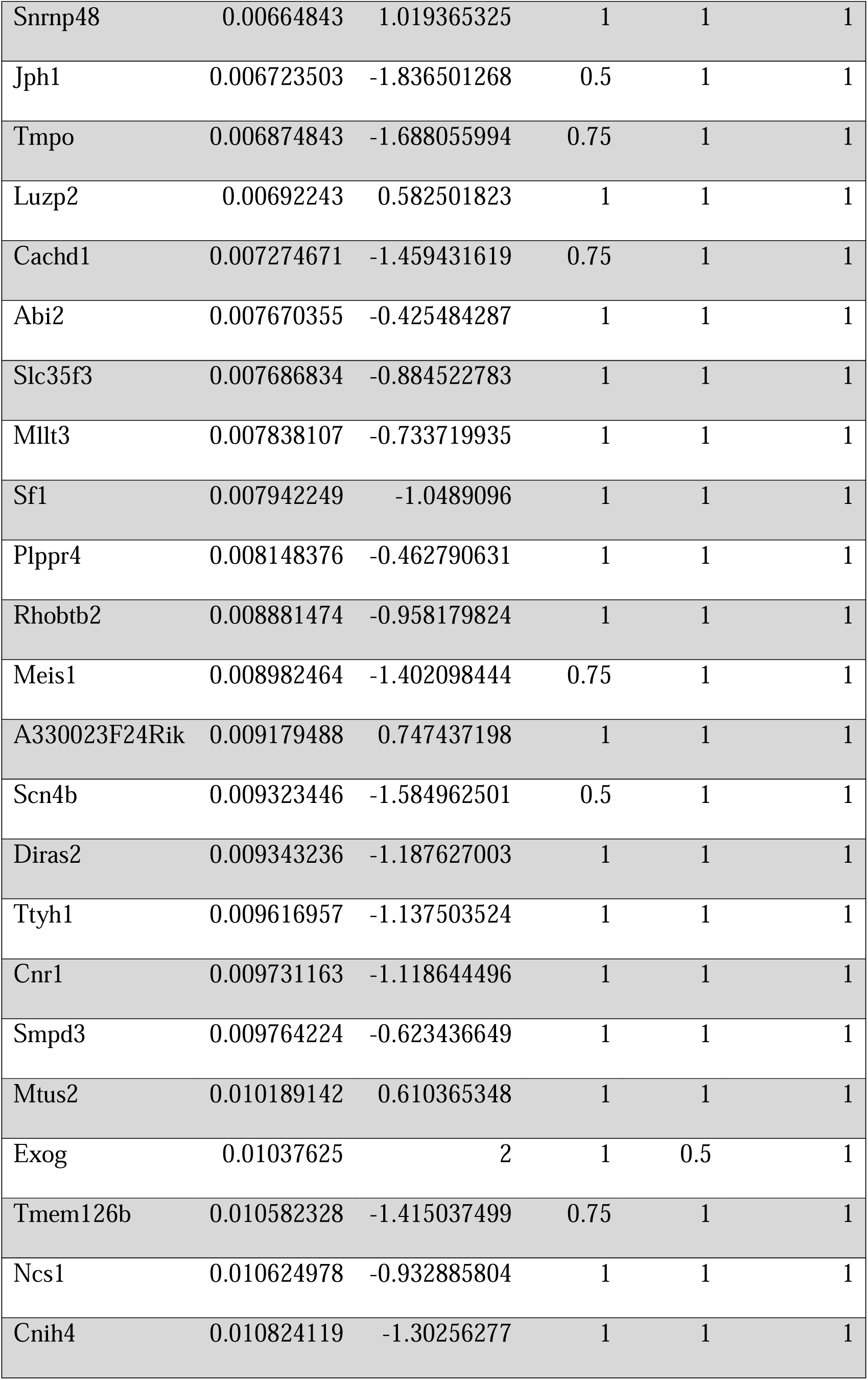
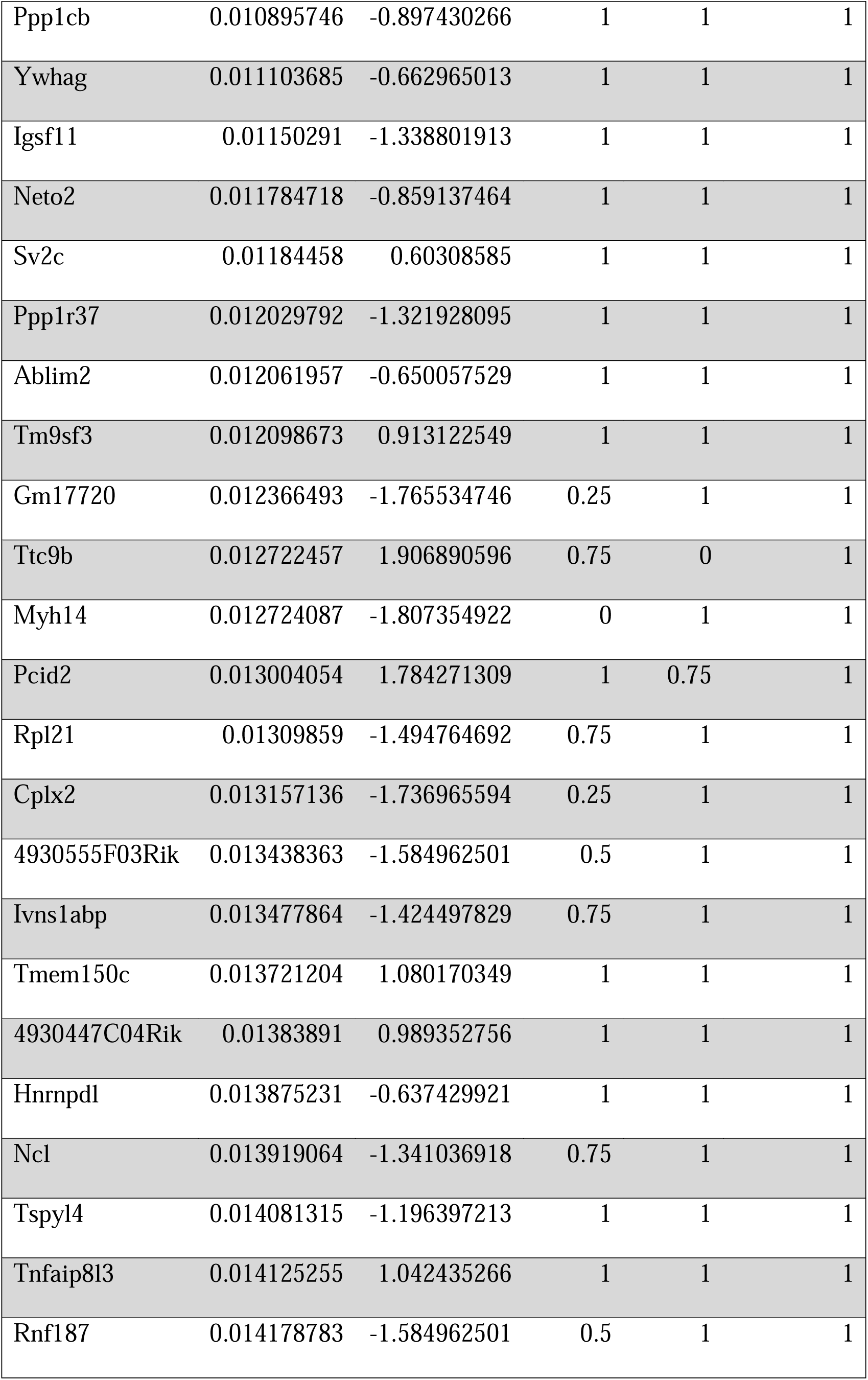
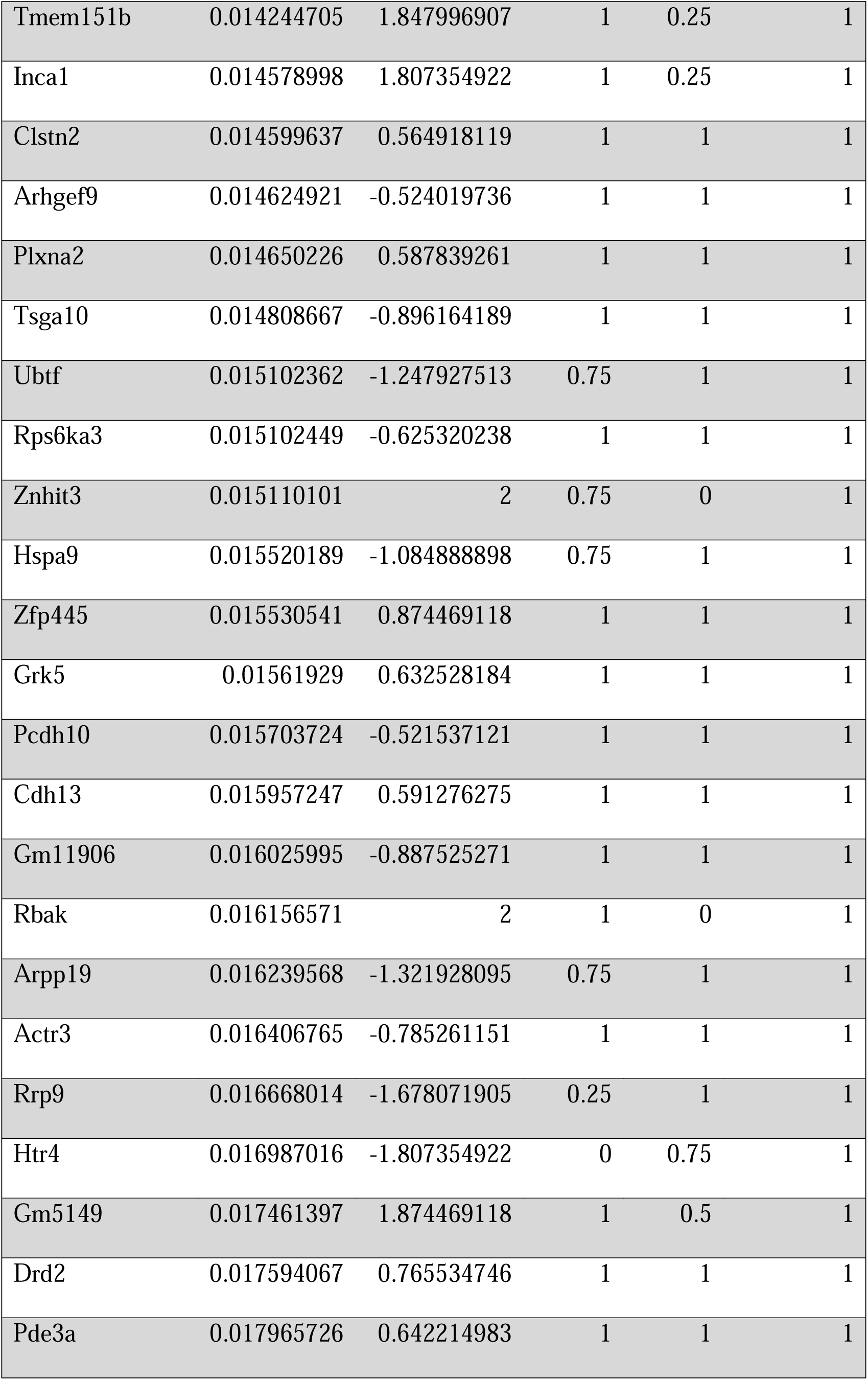
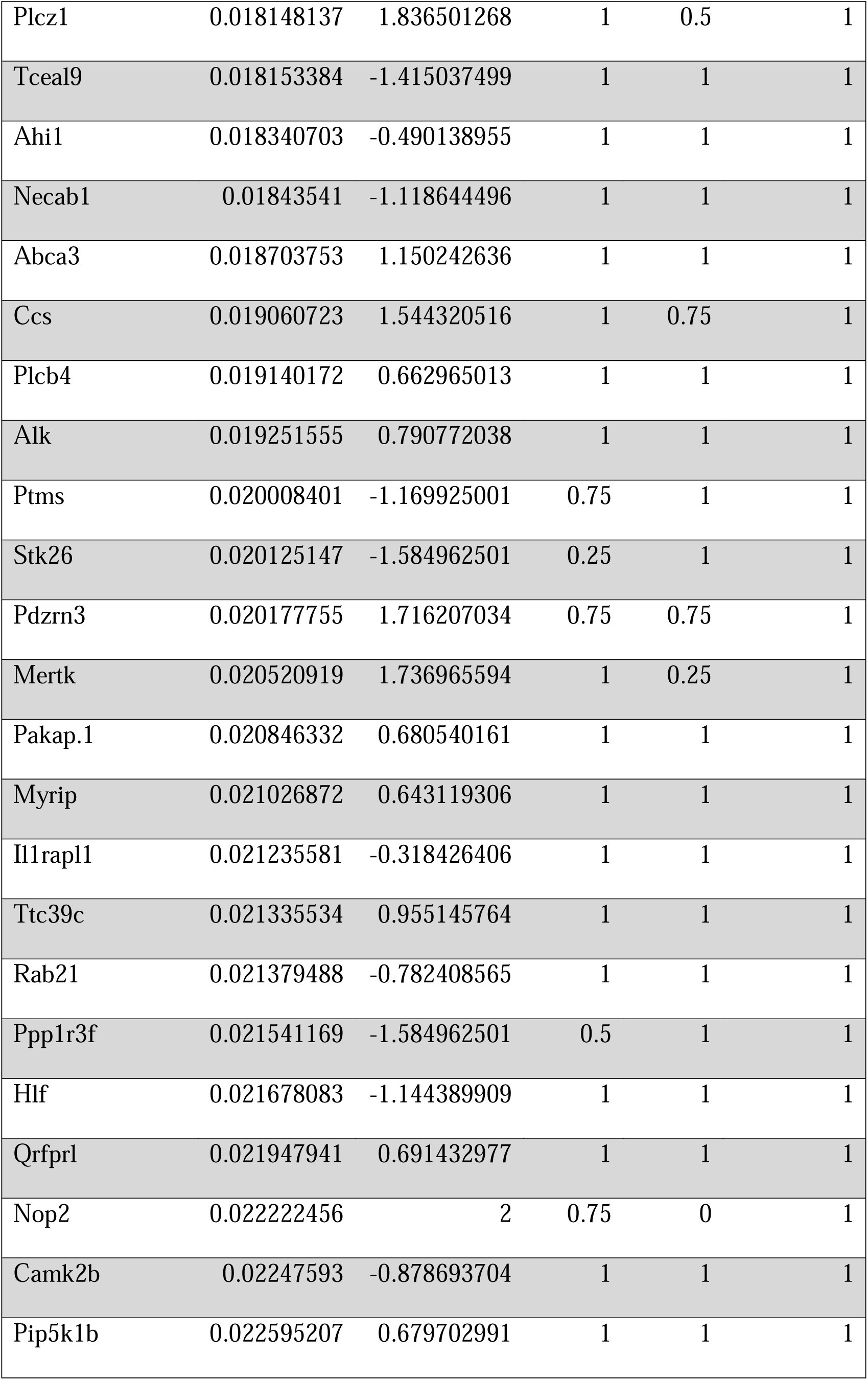
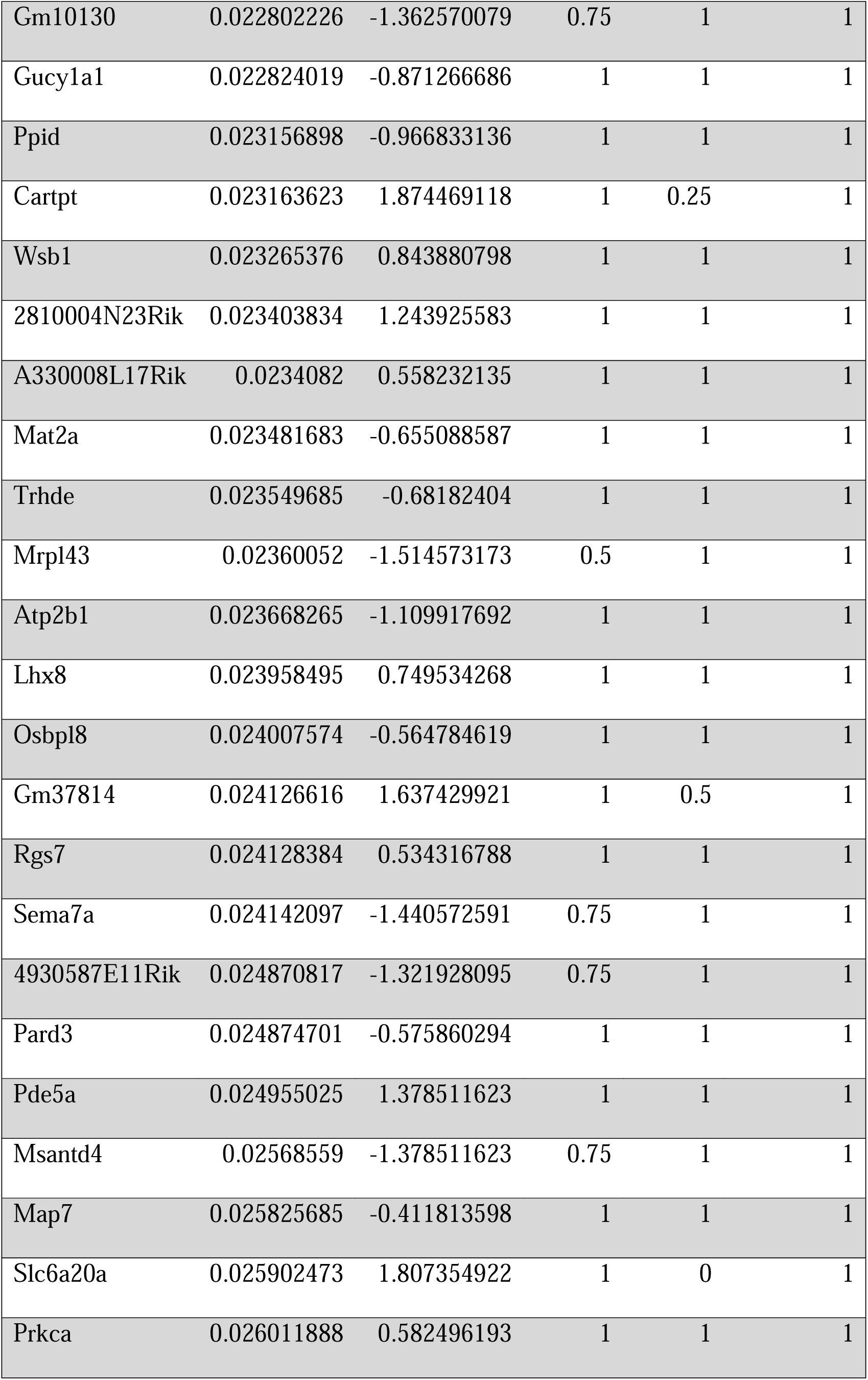
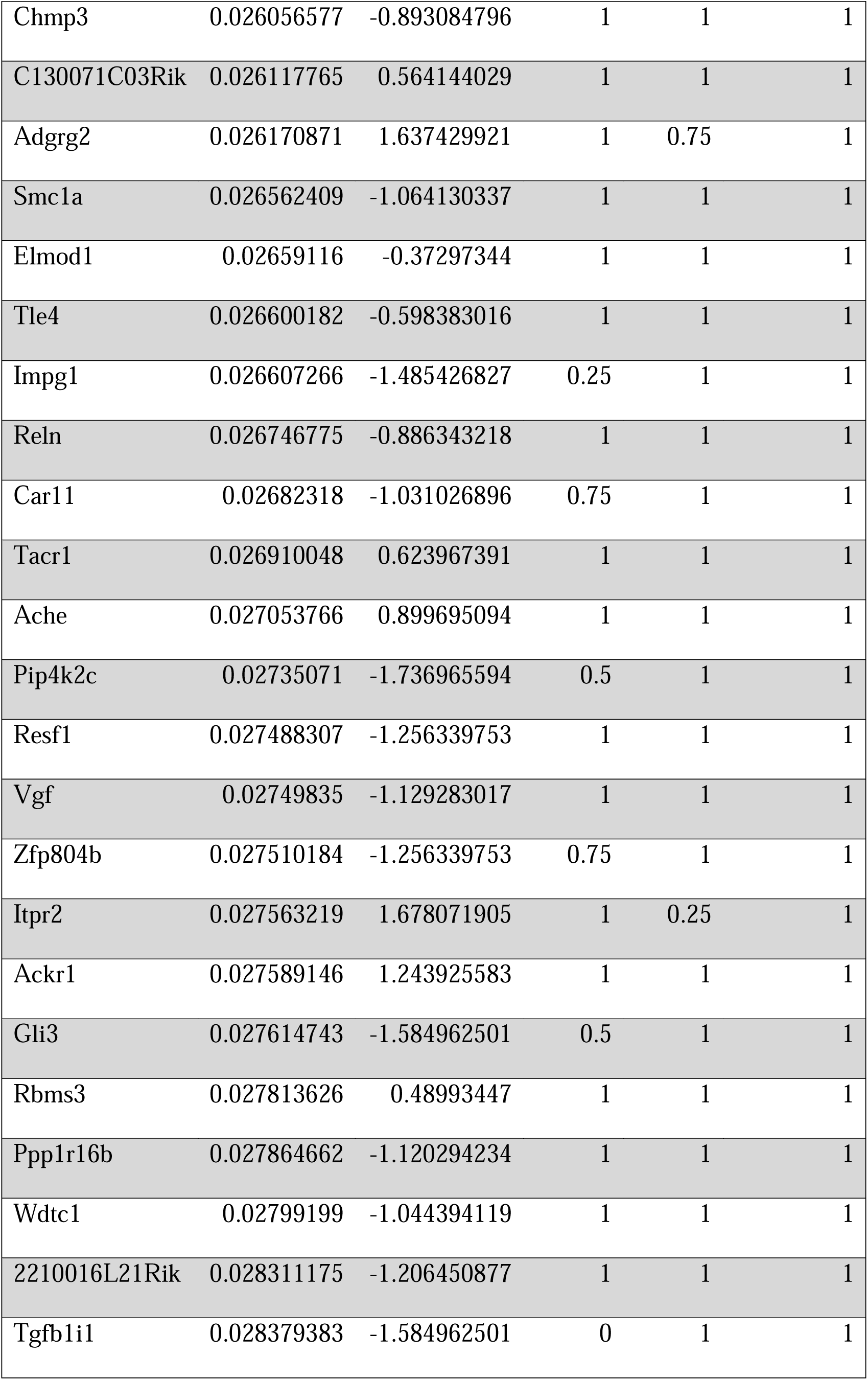
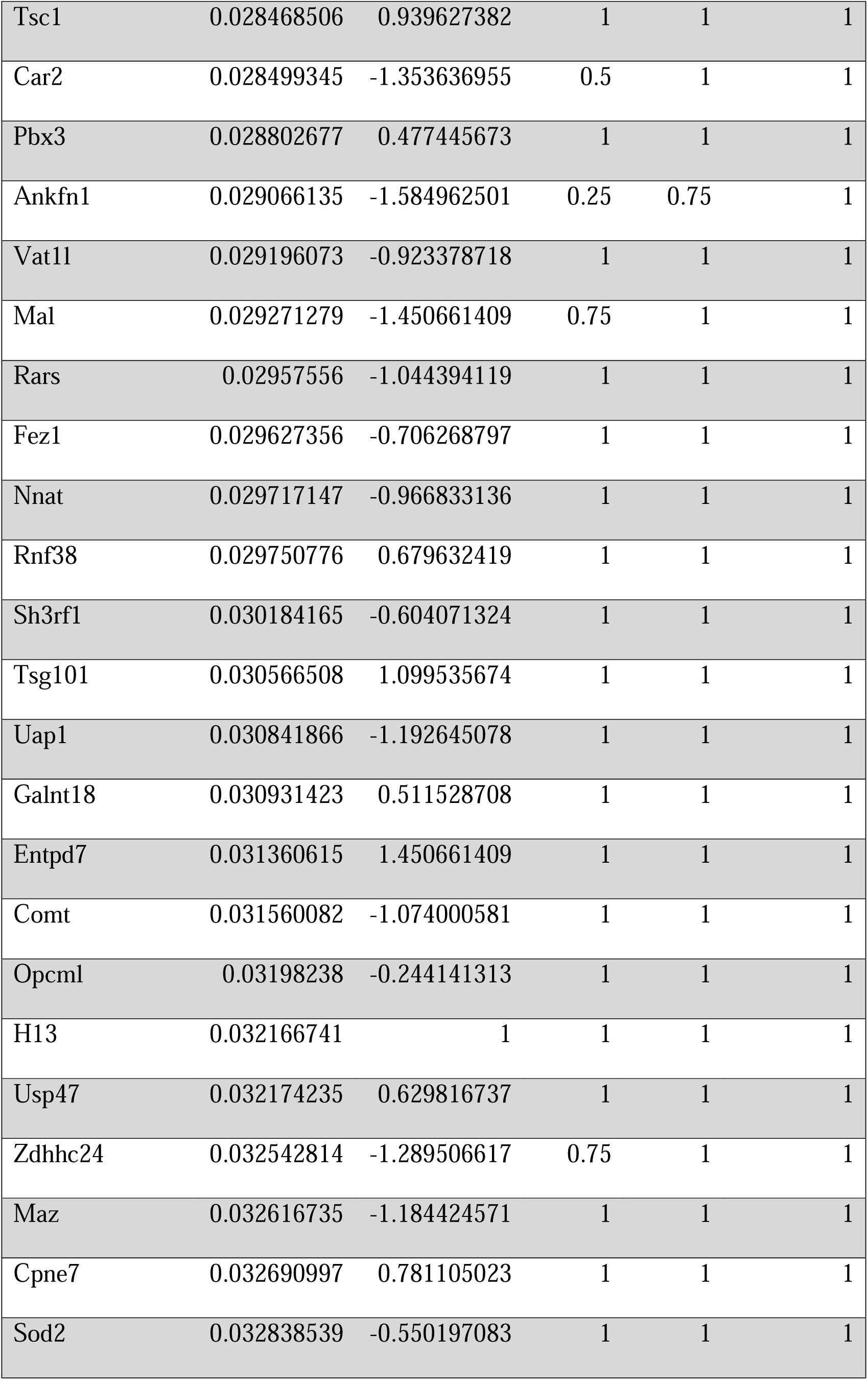
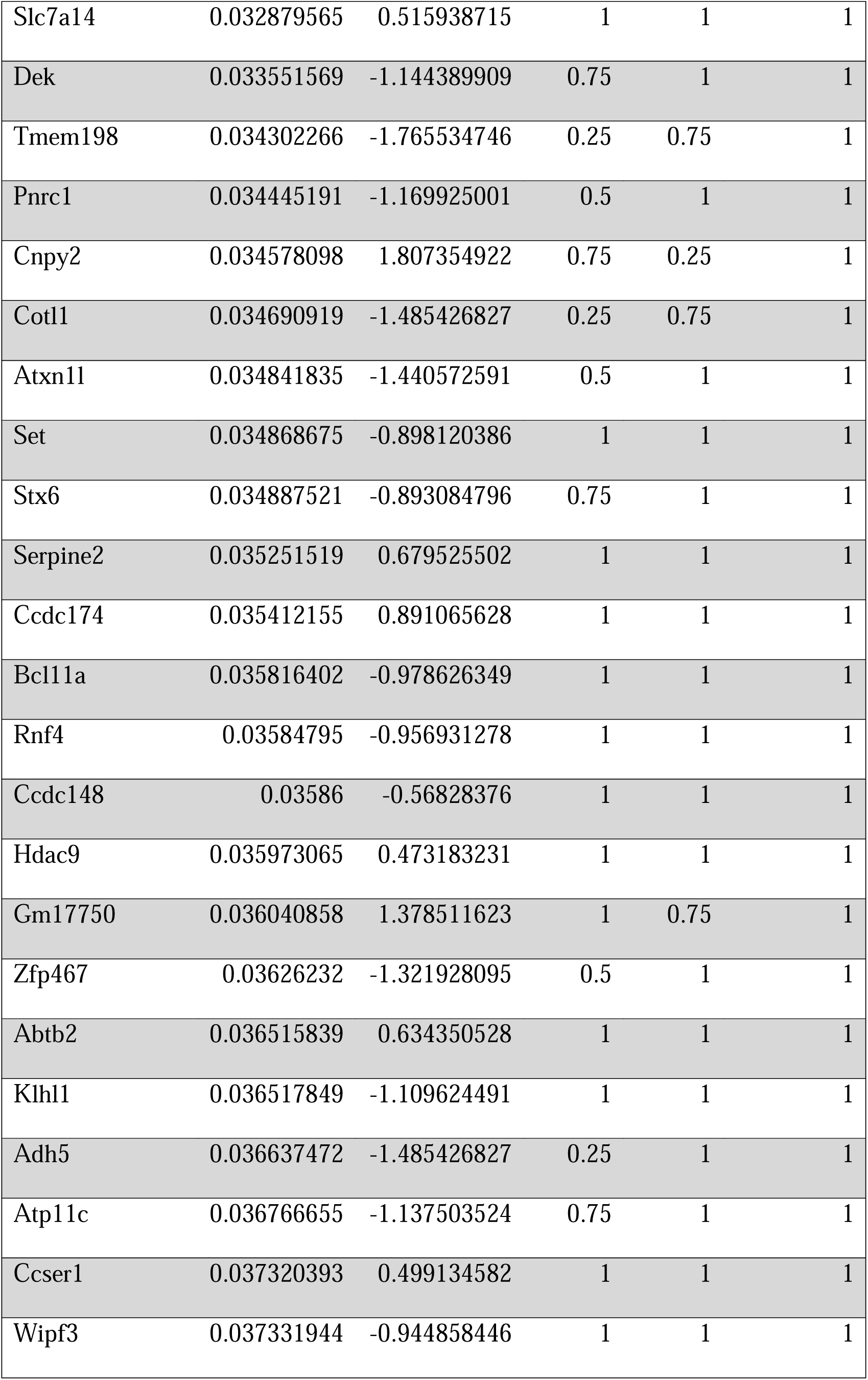
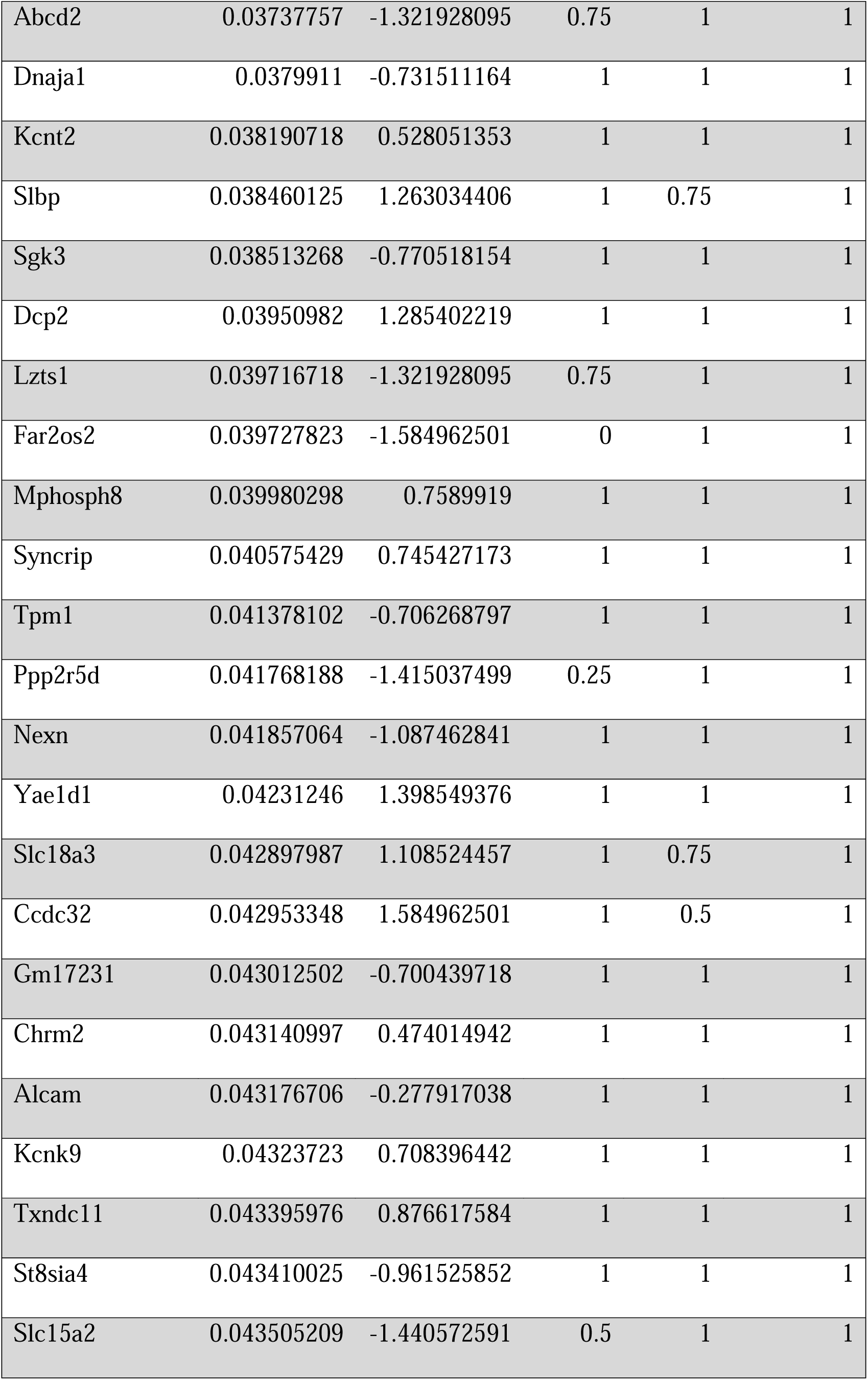
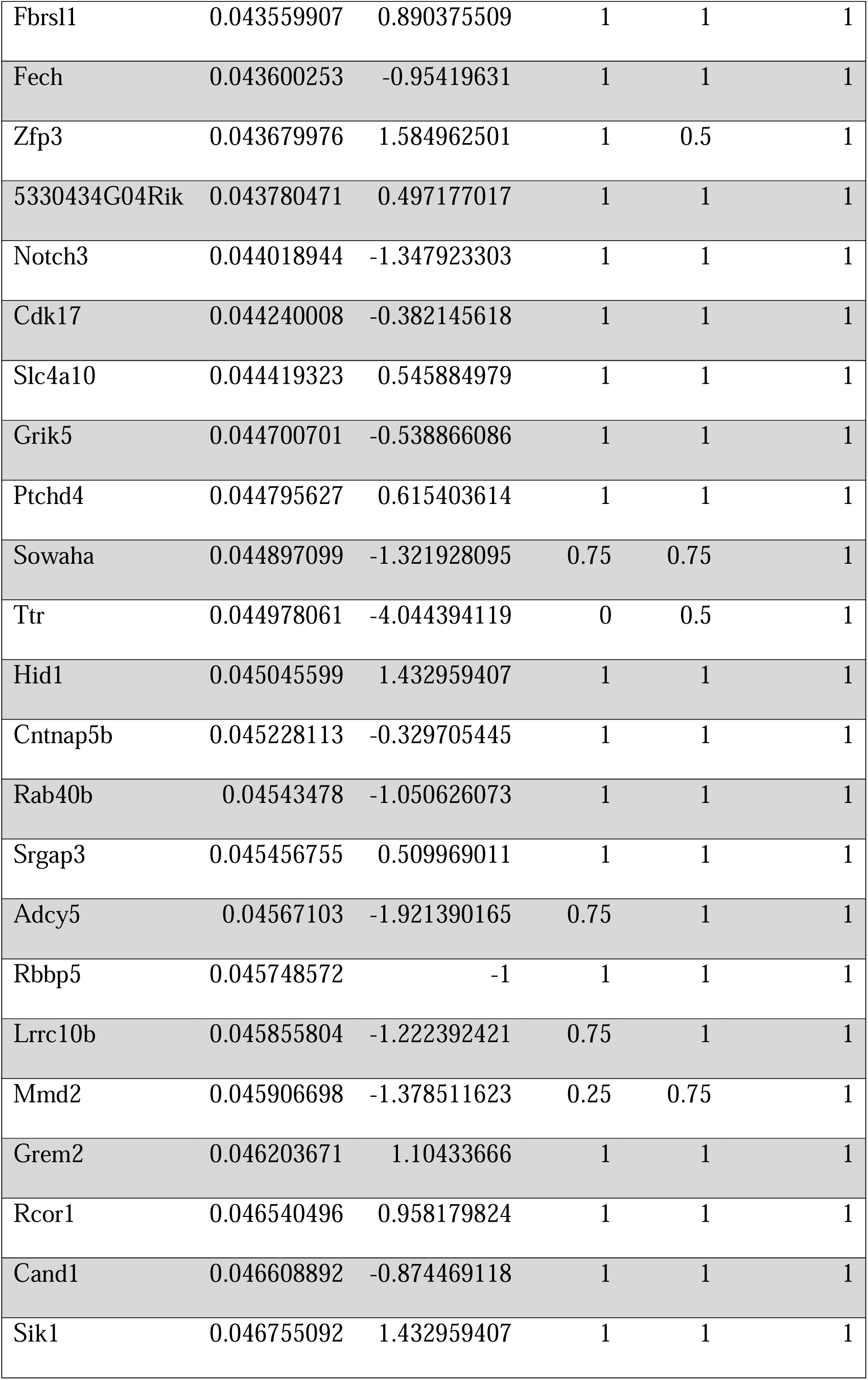
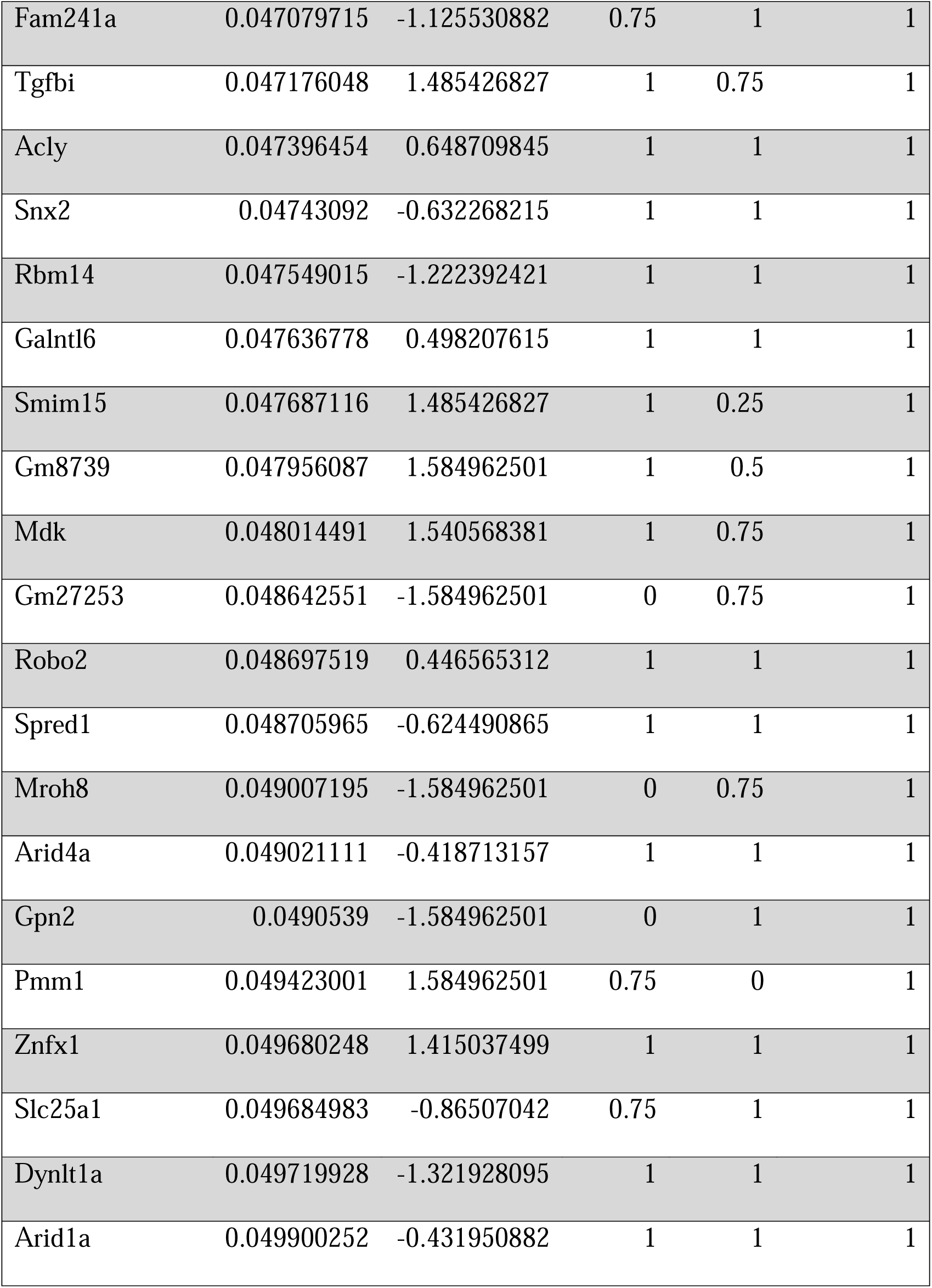
Differential expression results comparing *Df(16)1/+* and WT CHIs.

