## Supplementary figures and images for "Schizophrenia-associated 22q11.2 deletion elevates striatal acetylcholine and disrupts thalamostriatal projections to produce amotivation in mice"

### Figure S1

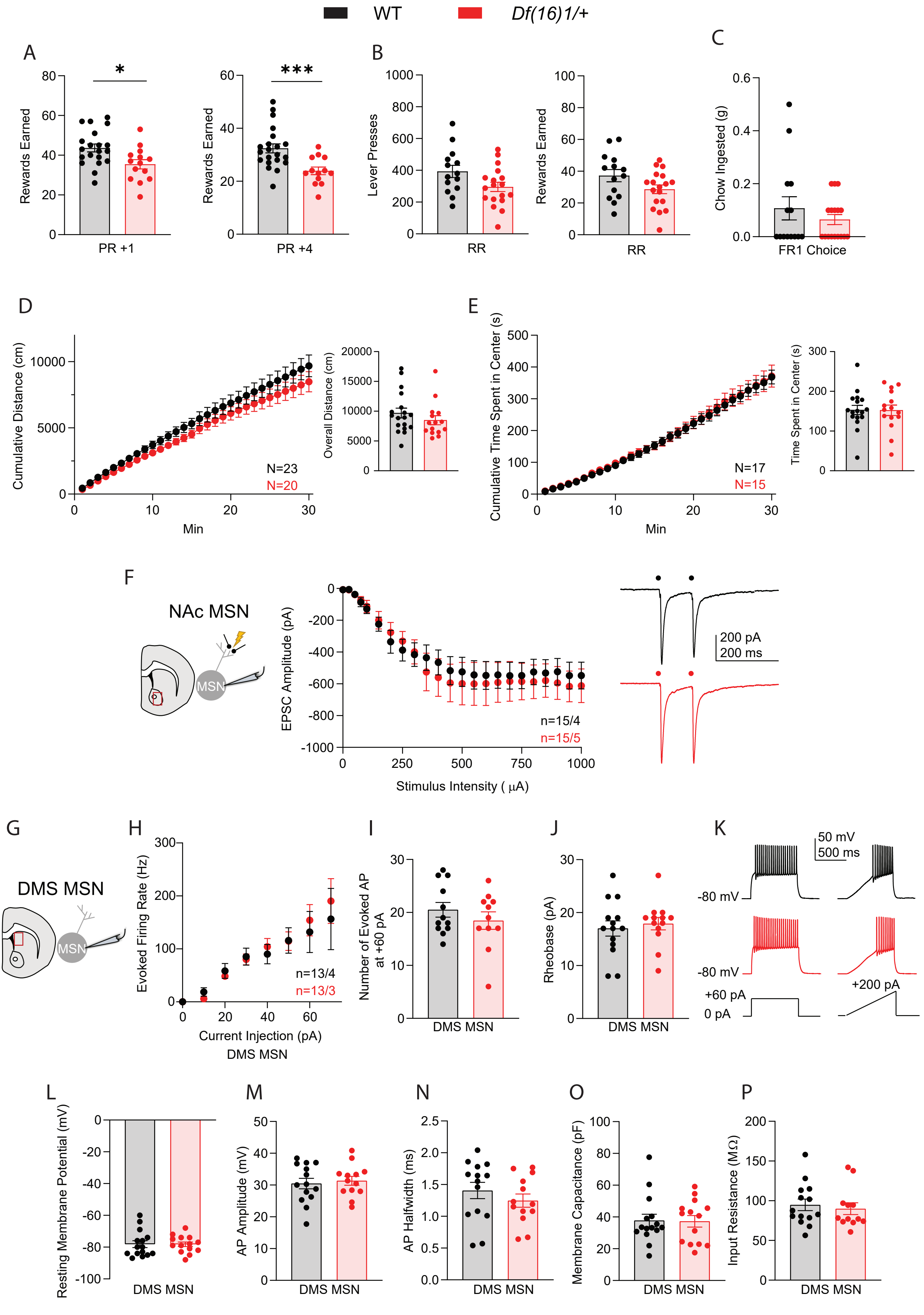

### Figure S2

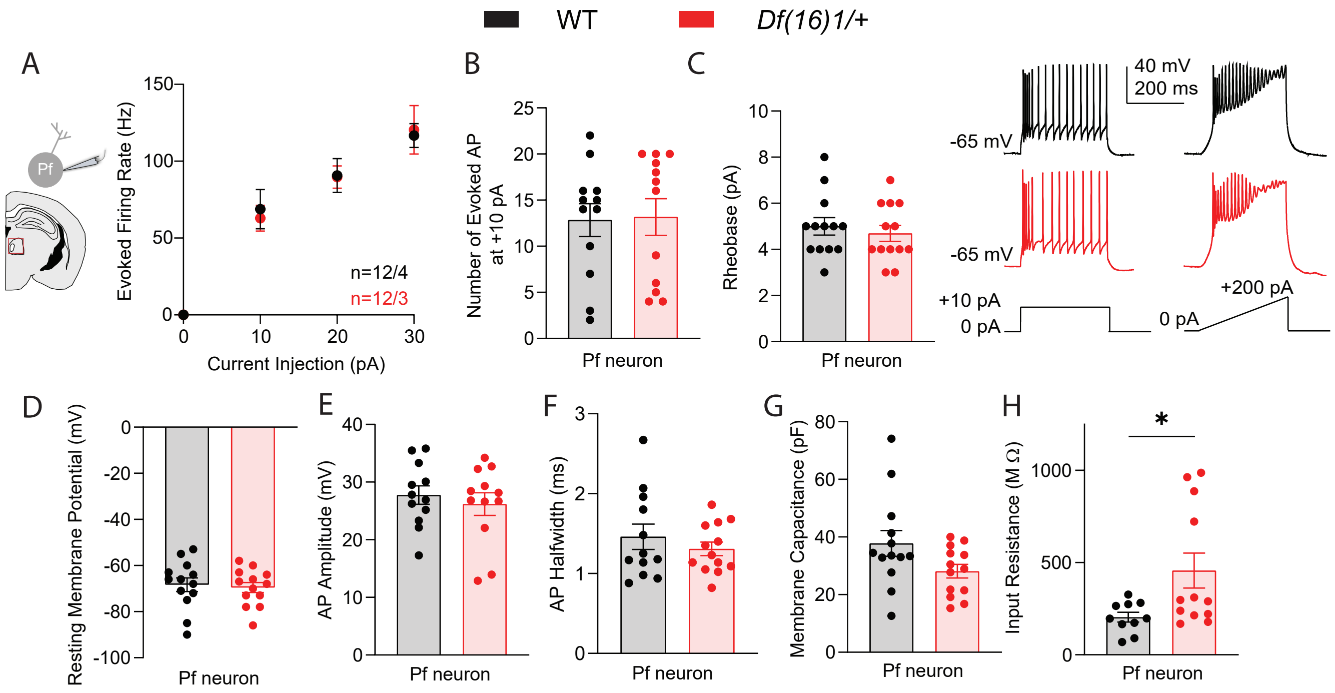

### Figure S3

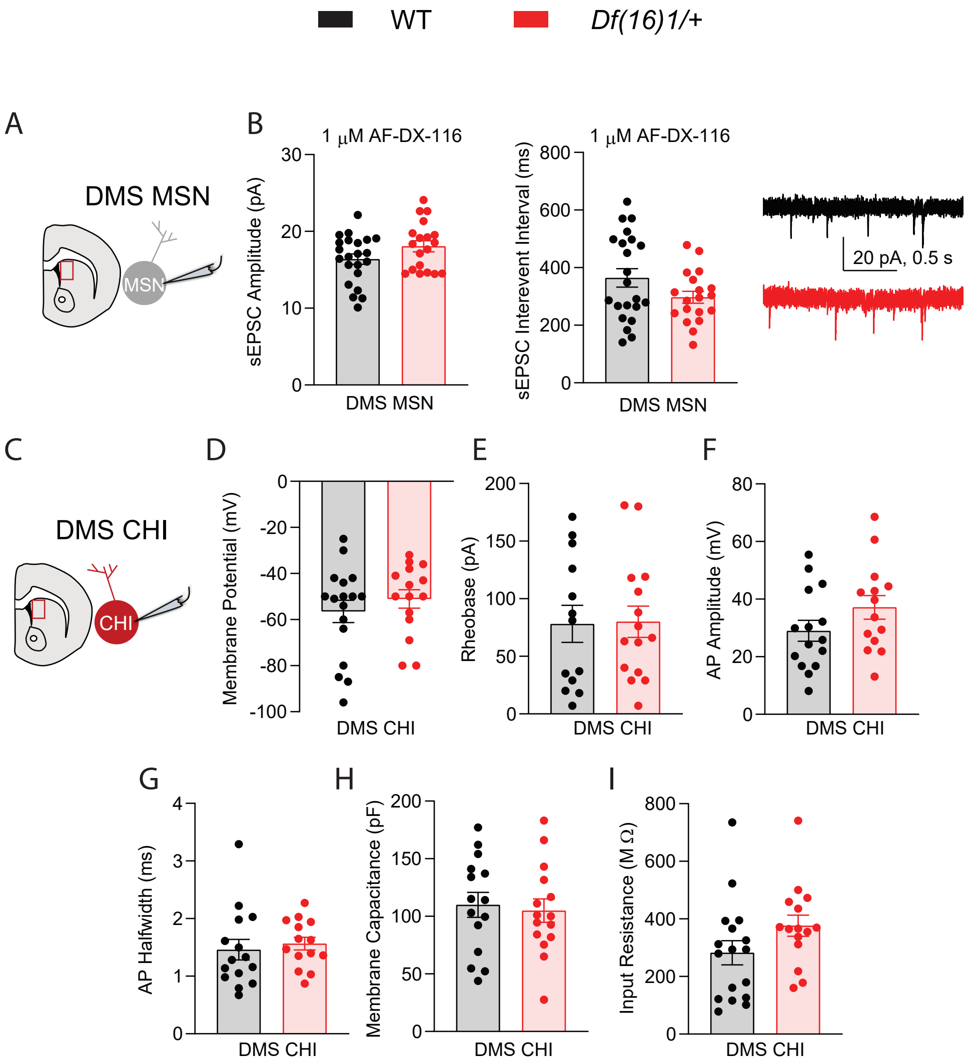

### Figure S4

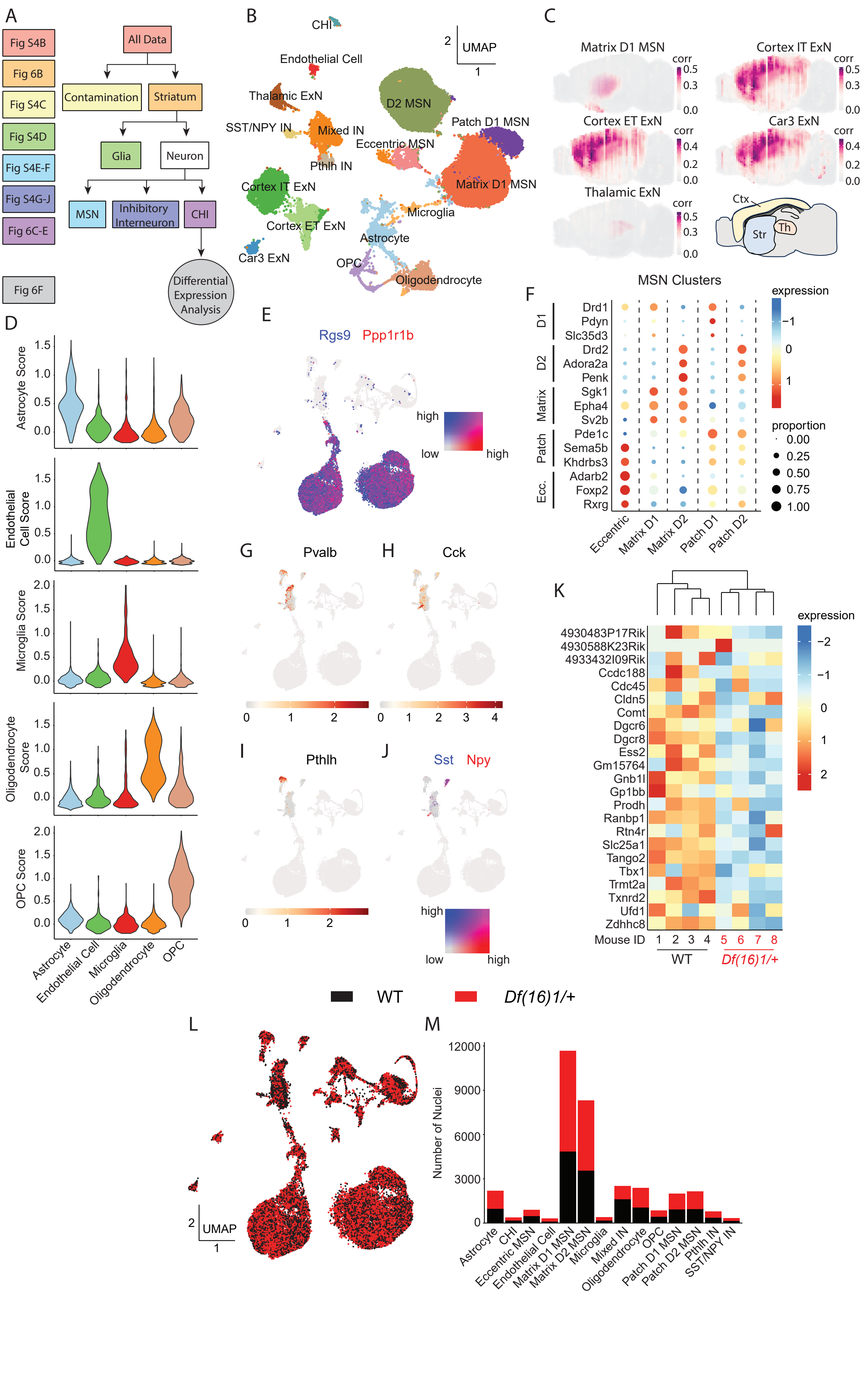

### Figure S5

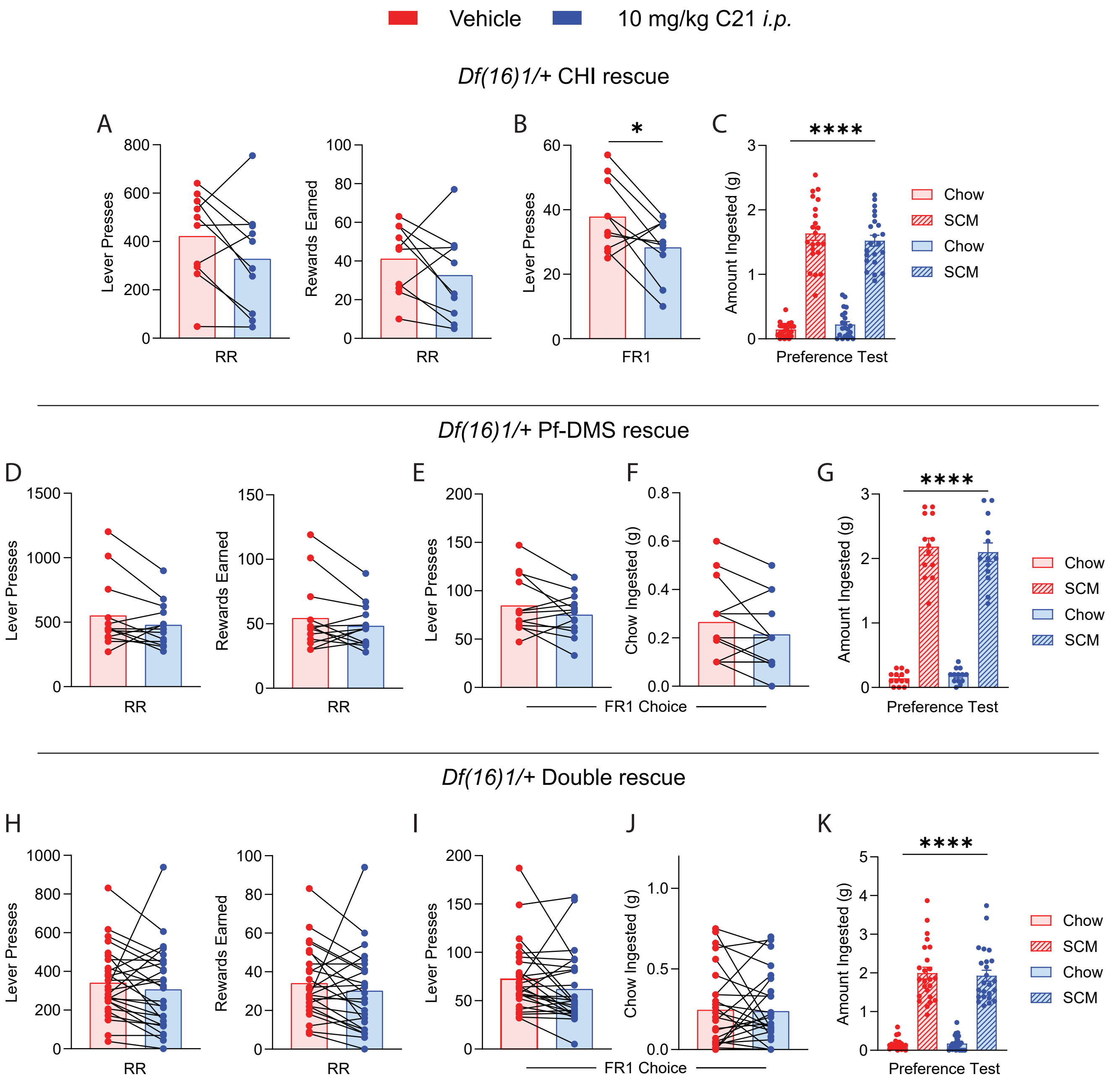
